# Regulation of proteasome activation by a ubiquitin-independent feedback mechanism

**DOI:** 10.64898/2026.06.05.730379

**Authors:** Samuel J. Watson, James C. Williamson, Pehuen P. Gerber, Guinevere L. Grice, Claudia Melucci, Aneisha Duggal, Christian M Gawden-Bone, Niek Wit, Richard T. Timms, James A. Nathan, Nicholas J. Matheson, Paul J. Lehner

## Abstract

The ubiquitin–proteasome system is the major pathway for selective protein degradation in eukaryotic cells. Some proteins are degraded by the proteasome without ubiquitination, but the prevalence and underlying mechanisms remain poorly understood. Here, we use pulsed-SILAC proteomics to systematically identify proteins undergoing ubiquitin-independent proteasomal degradation (*UbInPD*). We identify the paralogous proteasome activation factors ZFAND5 and ZFAND6 (ZFAND5/6) and show that ZFAND5 contains a ubiquitin-independent degron that can both promote its rapid turnover and allosterically activate the proteasome. These findings support a model in which proteasome activation and activator degradation are coupled through a ubiquitin-independent feedback mechanism. We further show that ZFAND5/6, together with the scaffold protein p62, restrain NF-κB activation in the TLR4 pathway. We link this regulation to the degradation of UBCH5c, an abundant E2 ubiquitin-conjugating enzyme that can initiate or ‘prime’ ubiquitin chain synthesis. Our findings expand the known landscape of *UbInPD* and reveal unexpected links between proteasome activation, E2 enzyme regulation, and inflammatory signalling.

## Introduction

All cells continuously identify and eliminate damaged, misfolded, or redundant proteins, making protein homeostasis (proteostasis) a fundamental cellular behaviour. Many debilitating diseases are caused by failures of proteostasis, and the loss of proteostasis is a hallmark of aging (Lopez-Otin *et al*, 2023).

The ubiquitin-proteasome system is the major pathway for selective protein degradation in eukaryotic cells, and it is implicated in essentially all cellular processes (Komander & Rape, 2012). Proteins destined for degradation are typically modified with ubiquitin via a three-enzyme cascade (E1 activating enzymes, E2 conjugating enzymes, E3 ligases) (Rape, 2018), and ubiquitin modification directs substrates to the 26S proteasome for degradation.

The 26S proteasome is a multi-subunit protease complex that unfolds and degrades ubiquitinated proteins (Collins & Goldberg, 2017). Protease activity is housed within the 20S core particle, while substrate recognition, unfolding, and entry are mediated by 19S regulatory particles that cap the 20S (Murata *et al*, 2009). The 19S regulatory particles contain receptors that selectively recognise ubiquitinated proteins (Collins & Goldberg, 2017). Some ubiquitinated proteins are also recruited to the proteasome via shuttle factors, which are ubiquitin receptors that recruit substrates to the proteasome but do not stably associate with it (Yu & Matouschek, 2017; Zientara-Rytter & Subramani, 2019).

The 26S proteasome’s proteolytic activity is allosterically regulated at steady state and following the accumulation of ubiquitinated proteins (Collins & Goldberg, 2017). Several factors activate the proteasome and enhance its ability to degrade its substrates. These include ZFAND5 (Collins & Goldberg, 2017), which is both a shuttle factor and a potent allosteric activator of the proteasome. ZFAND5’s activity regulates global protein turnover (Lee *et al*, 2018; Lee *et al*, 2023), and ZFAND5 has also been implicated in the regulation of inflammatory signalling (Hishiya *et al*, 2005; Huang *et al*, 2004).

Most proteins are targeted to the proteasome via ubiquitination. However, some proteins are directed to and degraded by the proteasome without ubiquitin modification (Erales & Coffino, 2014), and this phenomenon is known as ‘ubiquitin-independent proteasomal degradation’, or *UbInPD* (Makaros et al, 2023).

The classic example of a *UbInPD* protein is the metabolic enzyme ornithine decarboxylase (ODC), the rate-limiting enzyme in polyamine biosynthesis (Erales & Coffino, 2014). ODC is rapidly delivered to the proteasome via a dedicated, ubiquitin-independent targeting factor, Antizyme 1 (AZ1) (Murakami *et al*, 1992). AZ1 is rapidly induced when polyamines accumulate in cells, and it functions within a feedback circuit essential for cellular metabolism (Palanimurugan *et al*, 2004).

The transcription factor Rpn4 (Erales & Coffino, 2014) promotes the expression of proteasome subunits in budding yeast, and is also rapidly degraded by the proteasome via an intrinsic, ubiquitin-independent degron (Ha *et al*, 2012; Xie & Varshavsky, 2001). The rapid degradation of Rpn4 constitutes a feedback mechanism that ensures that proteasomes are produced at the correct rate, with Rpn4 acting as both a sensor and effector of proteasome production.

The recently described midnolin-proteasome pathway is a *UbInPD* pathway that promotes the degradation of various transcription factors (Gu *et al*, 2023). Midnolin acts like a shuttle factor by delivering proteins to the proteasome for degradation, but substrate recruitment and degradation are ubiquitin-independent (Nardone *et al*, 2025; Peddada *et al*, 2025).

While known *UbInPD* pathways regulate essential cellular processes, including metabolism, proteostasis, and cell signalling, the prevalence and underlying mechanisms of *UbInPD* remain incompletely understood. Here, we use pulsed-SILAC proteomics to systematically identify *UbInPD* proteins in human cells. We identify ZFAND5 and ZFAND6 as rapidly degraded *UbInPD* proteins, and show that they function within a ubiquitin-independent pathway that regulates proteasome activation. Furthermore, we identify a role for ZFAND5/6 in the regulation of NF-κB activation and inflammatory signalling. Proteomic experiments reveal that ZFAND5 associates with the scaffold protein p62, and that the two proteins convergently regulate the abundance of UBCH5c, an E2 ubiquitin-conjugating enzyme central to NF-κB activation.

## Results

### A pulsed-SILAC screen for *UbInPD* proteins

We designed a proteomic screen to systematically identify *UbInPD* proteins in HeLa cells (**Fig. 1A**). A pulsed-SILAC approach was chosen because this technique directly reports on protein stability (Ong *et al*, 2002). We reasoned that *UbInPD* proteins should be stabilised by treatment with a proteasome inhibitor (Bortezomib) but not by an inhibitor of global ubiquitination (TAK-243). To optimize treatment conditions, we determined the drug concentrations that maximally stabilise two unstable proteins targeted by known E3 ubiquitin ligases in HeLa cells (**Fig. S1A-C**), and confirmed that the optimised TAK-243 treatment markedly downregulates global ubiquitin conjugation (**Fig. S1D**).

**Figure 1.**
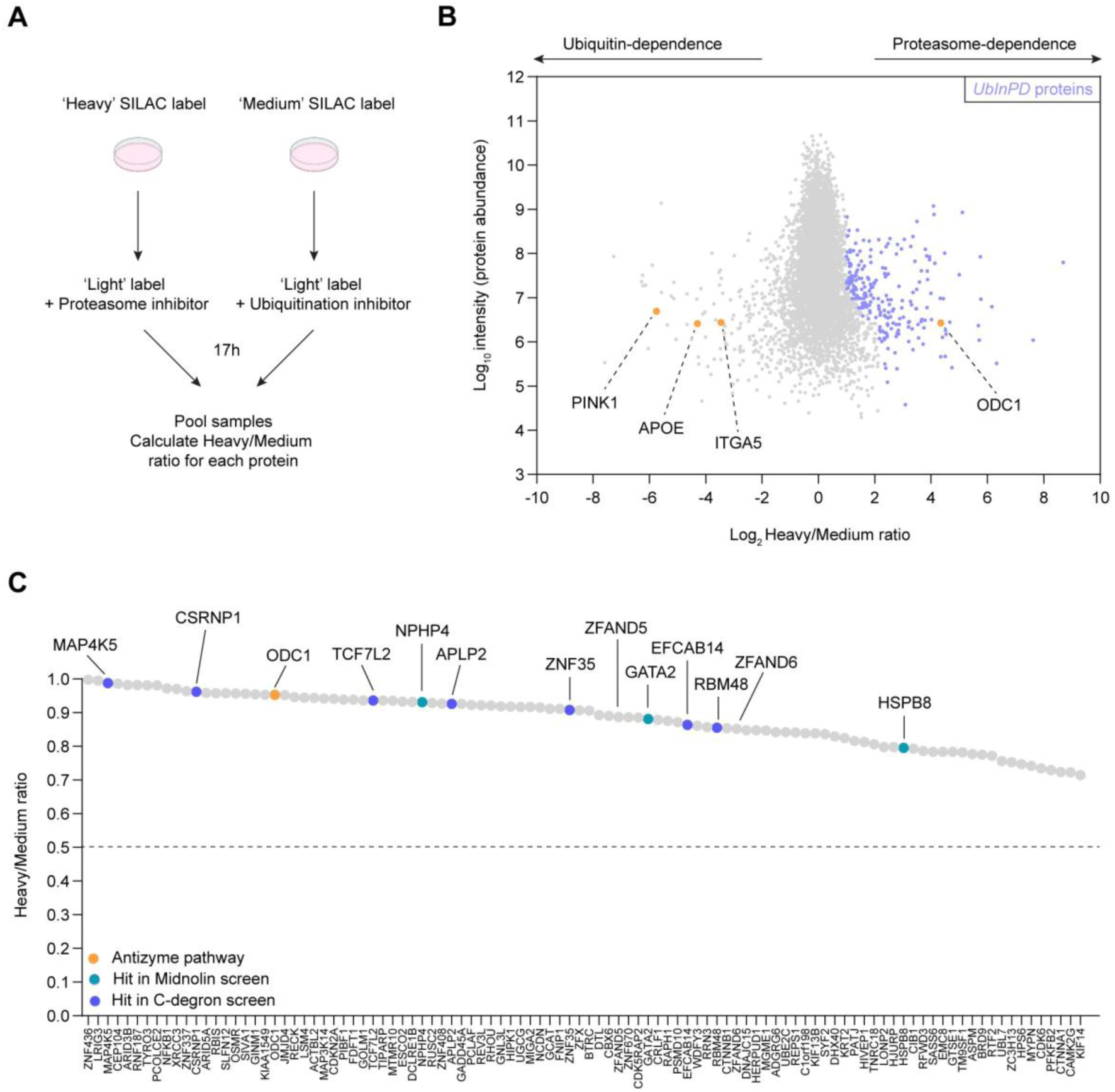
A pulsed-SILAC screen for *UbInPD* proteins. **(A)** Strategy to identify *UbInPD* proteins. Proteasome inhibitor: Bortezomib (100 nM). Ubiquitination inhibitor: TAK-243 (4.5 µM). SILAC: Stable isotopic labelling with amino acids in cell culture. *UbInPD*: Ubiquitin-independent proteasomal degradation. **(B)** Scatterplot displaying the results of the pulsed-SILAC proteomics. Log_2_ Heavy/Medium ratios are displayed on the X-axis, and overall protein abundance values are displayed on the Y-axis. Candidate *UbInPD* proteins are shown in purple and skew to the right because they are differentially stabilised by the proteasome inhibitor. **(C)** Plot displaying the highest confidence *UbInPD* proteins detected in the experiment. These proteins have a log_2_ Heavy/Medium ratio of >1 and a Significance B value of <0.001. The hits were cross-referenced against a screen for Midnolin substrates and a screen for *UbInPD* C-degrons (Gu *et al*., 2023; Makaros *et al*., 2023). Any H/M ratio above the dashed line is positive; any ratio below it is negative.

For the pulsed-SILAC screen, HeLa cells labelled with either ‘Heavy’ or ‘Medium’ amino acids were transferred to ‘Light’ media and treated with either Bortezomib or TAK-243. Cells were pooled after treatment and Heavy/Medium (H/M) ratios were calculated for each detected protein. We predicted that *UbInPD* proteins would be stabilised by Bortezomib but not TAK-243, resulting in positive H/M ratios, whereas proteins degraded through ubiquitin-dependent but proteasome-independent pathways would show negative H/M ratios.

We defined putative *UbInPD* proteins using the following criteria: (i) a log_2_ H/M ratio of >1, and (ii) a Significance B value of <0.05. Of the 7036 proteins detected within the experiment, 231 proteins satisfied these criteria. ODC, the classic positive control protein, was among the top hits, validating our approach (**Fig. 1B**). We also identified proteins whose turnover is ubiquitin, but not proteasome, dependent. These included plasma membrane proteins (e.g. ITGA5), extracellular proteins (e.g. APOE), and mitochondrial proteins (e.g. PINK1), which can be routed to the lysosome for degradation in a ubiquitin-dependent manner (**Fig. 1B**). Cross-referencing our gene list against a genetic screen for Midnolin substrates identified 8 overlapping hits, including NPHP4, GATA2, and HSPB8 (Gu *et al*., 2023) (**Fig. S2A, B; Fig. 1C**). A recent genetic screen used a ’GPS-peptidome’ library encoding the C-terminal 23 residues of all human proteins fused to an N-terminal stability tag (Makaros *et al*., 2023). to identify *UbInPD* proteins. Comparing this dataset with our gene list revealed 16 overlapping hits, including MAP4K5, CSRNP1, ODC, TCF7L2, APLP2, ZNF35, EFCAB14, and RBM48 (**Fig. S2C, D; Fig. 1C**). Thus, our proteomic screen identified both known and novel *UbInPD* proteins.

### ZFAND5 and ZFAND6 are *UbInPD* proteins

Two of the highest confidence hits, ZFAND5 and the closely related ZFAND6 (ZFAND5/6), were further characterised as they are both extremely unstable and can directly associate with the proteasome (Lee *et al*., 2018; Li *et al*, 2021). ZFAND5/6 both had high H/M ratios in the pulsed-SILAC screen (**Fig. 2A**) and were stabilised by Bortezomib but not TAK-243 in a validation experiment (**Fig. 2B**). Strikingly, TAK-243 treatment strongly destabilised both proteins. This may reflect a competition effect between ZFAND5/6 and ubiquitinated proteins – the global depletion of ubiquitinated proteins may ‘free up’ proteasomes to degrade *UbInPD* proteins more rapidly (Makaros *et al*., 2023). ZFAND5/6 also behaved as *UbInPD* proteins in both cycloheximide chase (**Fig. 2C**) and flow-cytometry-based protein stability assays (**Fig. S3A**). TAK-243 inhibits both E1 ubiquitin activating enzymes with similar efficiency (Hyer *et al*, 2018). Nevertheless, we used CRISPR-Cas9 to delete the minor E1 UBA6 and confirmed that there was no effect on ZFAND5/6 abundance (**Fig. S4A**). Together, these results indicate that ZFAND5 and ZFAND6 are bona fide *UbInPD* proteins.

**Figure 2.**
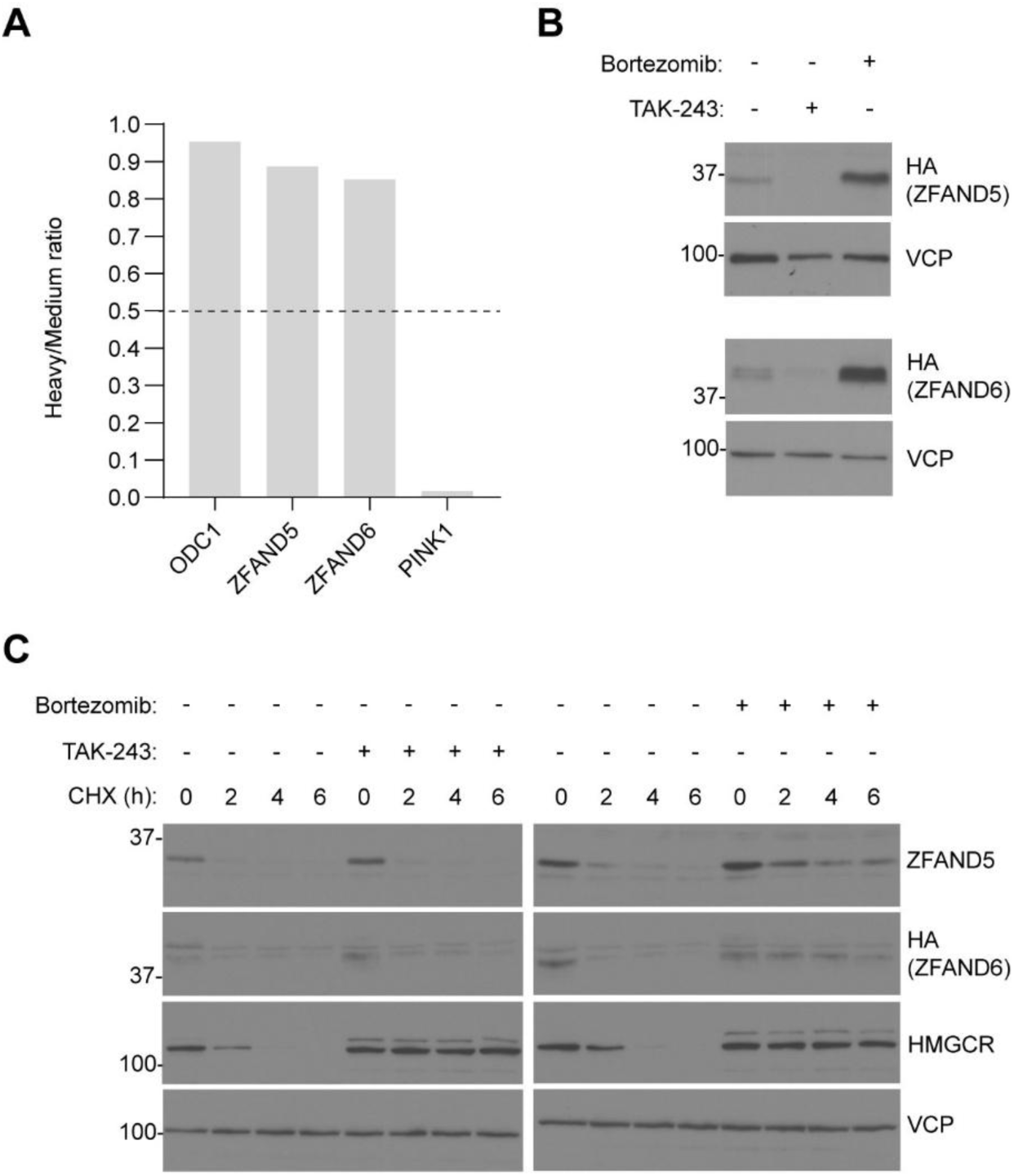
ZFAND5 and ZFAND6 are *UbInPD* proteins. **(A)** ZFAND5 and ZFAND6 (ZFAND5/6) were hits in the pulsed-SILAC proteomic screen for *UbInPD* proteins. ODC1 is included as a positive control. PINK1 is included as an example of a protein that is degraded by ubiquitin-dependent but proteasome-independent degradation. Any H/M ratio above the dashed line is positive; any ratio below it is negative. **(B)** Validation of ZFAND5/6 by immunoblotting. Cells stably overexpressing 3xHA ZFAND5 or ZFAND6 were treated with TAK-243 or Bortezomib overnight. Numbers to the left: molecular weight (kDa) here and in all subsequent immunoblots. **(C)** Validation of ZFAND5/6 by cycloheximide chase assay. Cells were treated with cycloheximide (10 µg/ml) across a 6-hour time-course. The cells stably overexpressed 3xHA ZFAND6. HMGCR: Control protein with known E3 ligases in HeLa cells. CHX: cycloheximide.

### ZFAND5 is degraded by a ubiquitin-independent, ‘proteasome activation degron’

Next, we sought to identify the ZFAND5/6 degron – the minimal region required for proteasomal degradation. ZFAND5 and ZFAND6 have an identical domain organisation, with three conserved domains (**Fig. 3A**). The A20 domain recruits ubiquitinated substrates, while the AN1 and CTR domains bind and allosterically activate the proteasome (Lee *et al*., 2023). Cycloheximide chase assays showed that ZFAND5/6 degradation depends on the AN1 domain, but not the A20 domain (**Fig. 3A**). Furthermore, A20 domain mutants remained *UbInPD* proteins, while AN1 mutants were completely stable (**Fig. S5A, B; Fig. S6A, B**).

**Figure 3.**
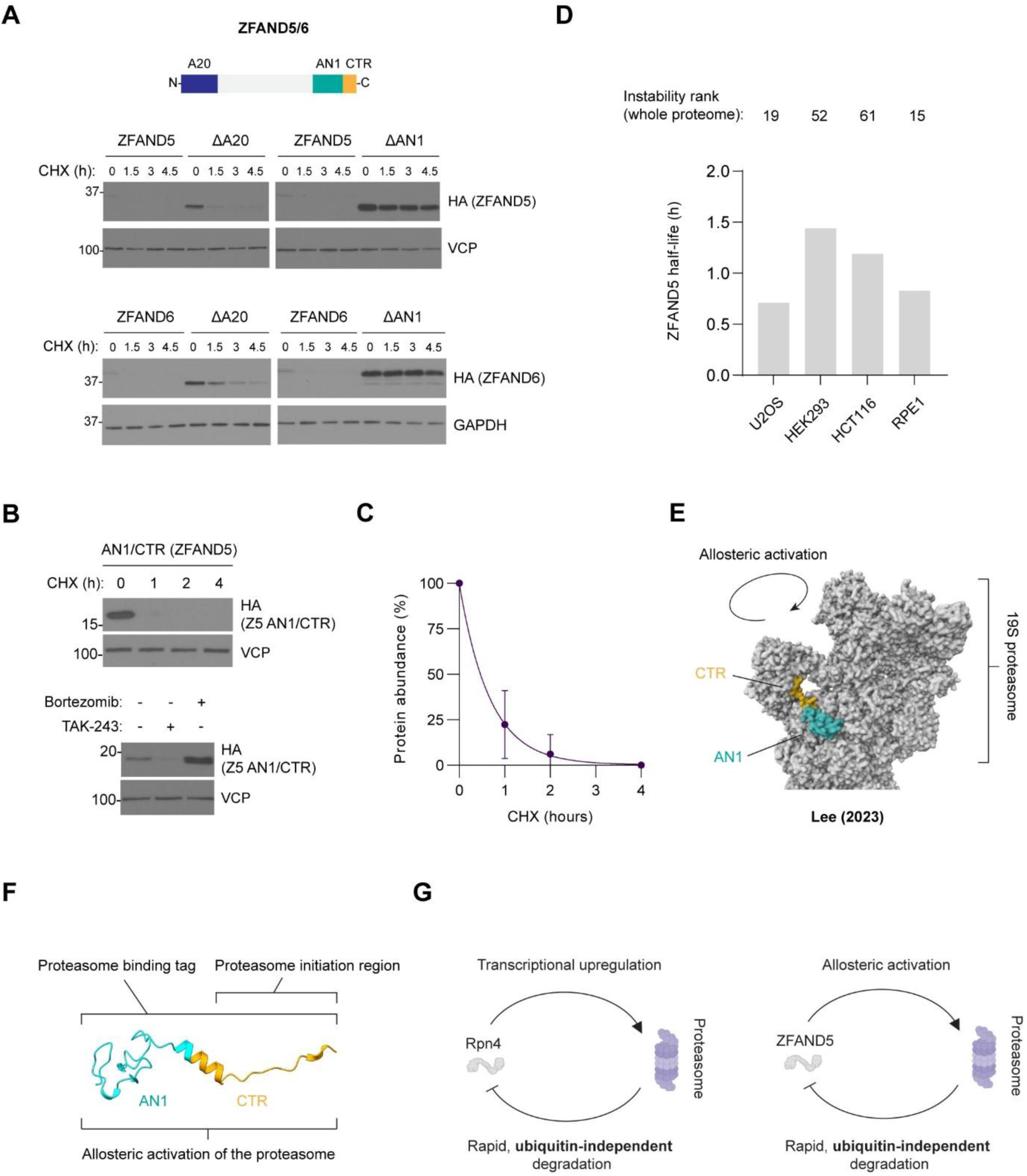
ZFAND5 is degraded by a ubiquitin-independent, ‘proteasome activation degron’. **(A)** ZFAND5/6 are rapidly degraded, and this degradation is dependent on the AN1 domain. ZFAND5/6 have an identical domain organisation, and this is shown at the top. Cells stably overexpressing ZFAND5/6 or the indicated truncation mutants (denoted by the Δ symbol) were treated with cycloheximide (10 µg/ml) across a 4.5-hour time-course. CHX: Cycloheximide. **(B)** The ZFAND5^AN1/CTR^ domain fragment is an unstable, *UbInPD* protein. Cells stably overexpressing 3xHA ZFAND5^AN1/CTR^ were treated with cycloheximide across the indicated time-course (above) or with TAK-243/Bortezomib for 17 hours (below). **(C)** Quantification of the cycloheximide chase experiment shown in (B). The experiment was performed three times, and the bands were quantified by densitometry. An exponential decay curve was fitted to the data. **(D)** Analysis of ZFAND5 stability in a publicly available dataset that profiled global protein stability across four cell lines (Li *et al*., 2021). ZFAND5 and ZFAND6 were part of a small set of proteins that were considered highly unstable across all four cell lines. ZFAND5’s half-life in each cell line is shown; numbers above the bars indicate the ranked instability of ZFAND5 (where 1 = the most unstable protein in the cell line). **(E)** Cryo-EM structure demonstrating the interaction between ZFAND5 and the proteasome (Lee *et al*., 2023). ZFAND5^AN1/CTR^ docks at a unique site centred around Rpt5, Rpt1, and Rpn1 and allosterically activates the proteasome as it engages. **(F)** Structure of ZFAND5’s ubiquitin-independent, ‘proteasome activation degron’. The degron contains elements that mediate proteasome binding and initiation and can also activate the proteasome upon engagement. **(G)** Model: Regulation of proteasome activation by a ubiquitin-independent feedback mechanism. Left: The Rpn4 transcription factor functions within a feedback circuit that regulates proteasome production in yeast. Right: ZFAND5 functions within a similar feedback circuit that regulates the allosteric activation of the proteasome in mammalian cells.

A ZFAND5 fragment consisting of just the AN1 and CTR domains remained a *UbInPD* protein and had an extremely short half-life (**Fig. 3B, 3C**). Both domains are highly conserved (**Fig. S7A-C**) and destabilised GFP when expressed as a GFP fusion protein (**Fig. S8A**). Thus, the AN1 and CTR domains constitute ZFAND5’s *UbInPD* degron. As ZFAND5 is among the most rapidly degraded proteins in the human proteome, our findings emphasise the potency of this degron (Li *et al*., 2021) (**Fig. 3D**). ZFAND5/6 degradation is unlikely to be a secondary consequence of proteasome engagement as other proteasome-binding proteins, including the canonical shuttle factor Rad23, are not degraded by the proteasome (Fishbain *et al*, 2011).

Degrons have two functional components: a ‘proteasome binding tag’ (usually a ubiquitinated lysine residue) and an unstructured ‘proteasome initiation region’ that is captured by the proteasome to initiate degradation (Inobe *et al*, 2011), and is usually N-or C-terminal (Fishbain *et al*., 2011). Recently published cryo-EM structures demonstrate that ZFAND5 binds to a unique site on the proteasome via its AN1/CTR domains and in so doing allosterically activates the proteasome (Lee *et al*., 2023) (**Fig. 3E**). These structures, together with our studies, explain how the AN1/CTR domains function as a ubiquitin-independent degron: The AN1/CTR domains form the ‘proteasome binding tag’, while the unstructured CTR domain likely acts as the ‘proteasome initiation region’ (**Fig. 3F**). The degron also has an additional feature – it contains elements sufficient to allosterically activate the proteasome (Lee *et al*., 2023). Together, these findings identify the ZFAND5 AN1/CTR region as a specialized ’proteasome activation degron’ that drives rapid, ubiquitin-independent turnover.

There are clear conceptual parallels between ZFAND5 and the Rpn4 transcription factor. In budding yeast, Rpn4 drives the expression of proteasome subunits and is also rapidly degraded by the proteasome via an intrinsic, ubiquitin-independent degron (Xie & Varshavsky, 2001). This feedback circuit ensures that proteasome production is titrated to cellular demand (**Fig. 3G**). We propose an analogous model for ZFAND5: by allosterically activating the proteasome while itself undergoing rapid, ubiquitin-independent degradation, ZFAND5 also acts as a self-limiting regulator (**Fig. 3G**). As ZFAND5 is an important regulator of global proteostasis (Lee *et al*., 2018), this feedback mechanism likely prevents excessive and deleterious activation of the proteasome.

### ZFAND5/6 restrain NF-κB activation in the TLR4 pathway

Although ZFAND5 has primarily been studied within the context of proteostasis, it is highly expressed in monocytes and macrophages, and conflicting reports suggest a role for ZFAND5 in the regulation of NF-κB signalling (Hishiya *et al*., 2005; Huang *et al*., 2004; Uhlen *et al*, 2015). A20/TNFAIP3, a classic negative regulator of NF-κB activation, contains an unusual ubiquitin binding domain (A20^ZnF4^) that is essential for NF-κB regulation *in vivo* (Martens *et al*, 2020). We noted that related domains in ZFAND5 and ZFAND6 (ZFAND5/6^A20^) (Enesa & Evans, 2014; Garner *et al*, 2011; Lee *et al*., 2023) are structurally similar to the A20^ZnF4^ domain (**Fig. 4A, B**) (Bosanac *et al*, 2010; Garner *et al*., 2011). These observations motivated us to investigate the association between ZFAND5/6 and inflammatory signalling.

**Figure 4.**
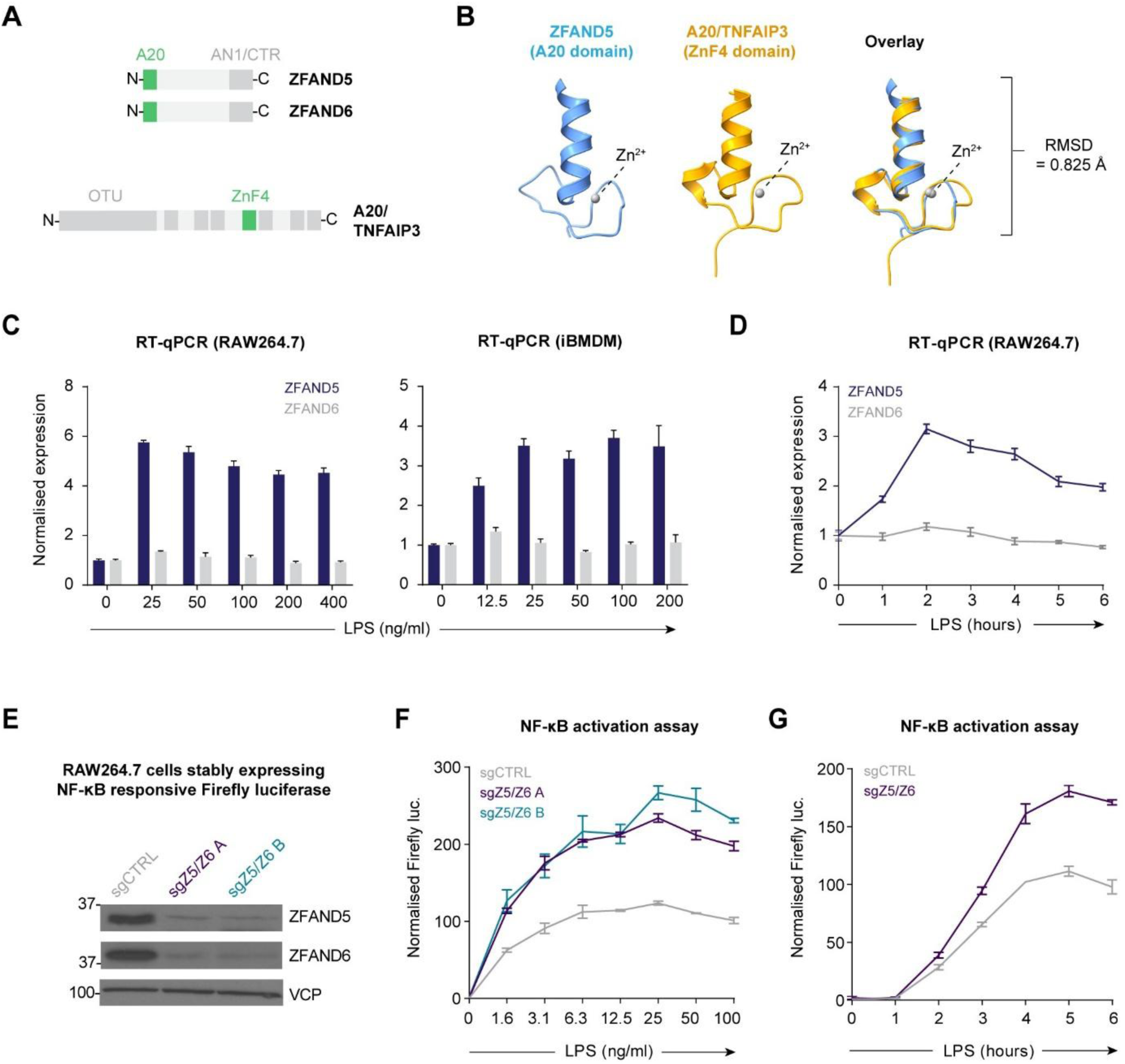
ZFAND5/6 restrain NF-κB activation in the TLR4 pathway. (A-B) The ZFAND5/6 A20 domain is closely related to A20/TNFAIP3’s ZnF4 domain. (A) ZFAND5, ZFAND6, and A20 domain maps. (B) Comparison of the ZFAND5^A20^ domain (PDB 2KZY) and the A20^ZnF4^ domain (PDB 3OJ3). RMSD: Root Mean Square Deviation. **(C-D)** Transcriptional induction of ZFAND5 in LPS-treated macrophages. Indicated cells (RAW264.7 cells and iBMDMs) were treated with an LPS titration for 24 hours (C) or with 10 ng/ml LPS across a 6-hour time-course (D). Expression was normalised to GAPDH and presented as fold change relative to the untreated control. Error bars: mean ± SD from three technical replicates. iBMDMs: Immortalised bone marrow-derived macrophages. **(E-G)** ZFAND5/6 restrain NF-κB activation. (E) Description of the NF-κB reporter cell line and CRISPR-Cas9-mediated deletion of ZFAND5/6. The reporter cells were treated with an LPS titration for 6 hours (F) or with 10 ng/ml LPS across a 6-hour time-course (G). Firefly signal was normalised against Renilla signal and presented as fold change relative to the untreated, non-targeting sgRNA control (sgCTRL). Error bars: mean ± SD from three technical replicates. Luc: luciferase.

As ZFAND5 is highly expressed in macrophages, we examined the expression of ZFAND5/6 in RAW264.7 cells (a mouse macrophage cell line) and in murine immortalised bone marrow-derived macrophages (iBMDMs) following LPS stimulation to activate NF-κB via the TLR4 pathway. ZFAND5 expression increased rapidly after LPS treatment, while ZFAND6 was constitutively expressed and insensitive to LPS (**Fig. 4C-D**; **Fig. S9A-B**). In addition, proteasome inhibition strongly stabilised ZFAND5, confirming its rapid degradation in mouse cells (**Fig. S9C**).

To determine whether ZFAND5/6 regulate NF-κB signalling, we used CRISPR-Cas9 to delete both genes individually and in combination in a dual-luciferase NF-κB reporter cell line (**Fig. S10A-D**). Because ZFAND5 and ZFAND6 are closely related, we analysed both genes to address possible functional redundancy. ZFAND5 knockout cells showed a marked increase in NF-κB reporter activity after LPS stimulation, whereas ZFAND6 knockout cells showed only a marginal increase (**Fig. S11A**). However, the ZFAND5/6 double knockout cells displayed a clear additive phenotype and the strongest overall increase in signalling. The enhanced signalling observed in the double knockout cells was consistent across a broad range of LPS concentrations and could be detected within the first two hours of stimulation (**Fig. 4E-G**).

To determine whether ZFAND5/6 regulate the expression of endogenous NF-κB target genes, we performed RNA-seq on control and ZFAND5/6 knockout cells treated with LPS across a short time-course (**Fig. S12A**). Consistent with the reporter assays, the NF-κB target genes *Tnf*, *Cxcl2* and *Tnfaip3* displayed enhanced induction in ZFAND5/6-deficient cells, a finding that was readily confirmed by RT-qPCR (**Fig. S12B-D**).

To explore the relationship between ZFAND5, ZFAND6, and A20, we compared ZFAND5/6 double knockout cells, A20 single knockout cells, and ZFAND5/6/A20 triple knockout cells (**Fig. S13A**). The ZFAND5/6 double knockout cells showed a stronger phenotype than the A20 knockout cells, but the triple mutant showed the strongest overall response. The additive phenotype observed in the triple knockout cells suggests that ZFAND5/6 and A20 regulate NF-κB via mechanisms that are at least partially independent. Taken together, our data indicate that ZFAND5 and ZFAND6 restrain NF-κB activation in the TLR4 pathway.

### ZFAND5/6 regulate the abundance of UBCH5c

Since ZFAND5/6 are shuttle factors that transport proteins to the proteasome for degradation, we hypothesised that ZFAND5/6 might restrain NF-κB signalling by preferentially promoting the degradation of important regulatory proteins. To identify such proteins, we used quantitative proteomics to compare the proteomes of control and ZFAND5/6 knockout cells, both with and without LPS stimulation. The E2 ubiquitin-conjugating enzyme UBCH5c was strongly upregulated in the ZFAND5/6 knockout cells at steady state and after LPS treatment, whereas UBC13 – an E2 with a well-established role in NF-κB signalling – was unaffected (**Fig. 5A**; **Fig. S14A**). Additionally, there was no change in UBCH5c mRNA abundance between the samples, indicating post-transcriptional regulation (**Fig. 5B**). UBCH5c is an abundant and promiscuous E2 ubiquitin-conjugating enzyme that functions with multiple E3 ligases, including several that drive NF-κB activation (Roman-Trufero & Dillon, 2022; Shembade *et al*, 2010). Notably, A20 interacts with UBCH5c and UBC13 through its A20^ZnF4^ domain, and it promotes the turnover of both E2s (Bosanac *et al*., 2010; Shembade *et al*., 2010). These observations suggested that ZFAND5/6 might restrain NF-κB signalling by regulating UBCH5c abundance.

**Figure 5.**
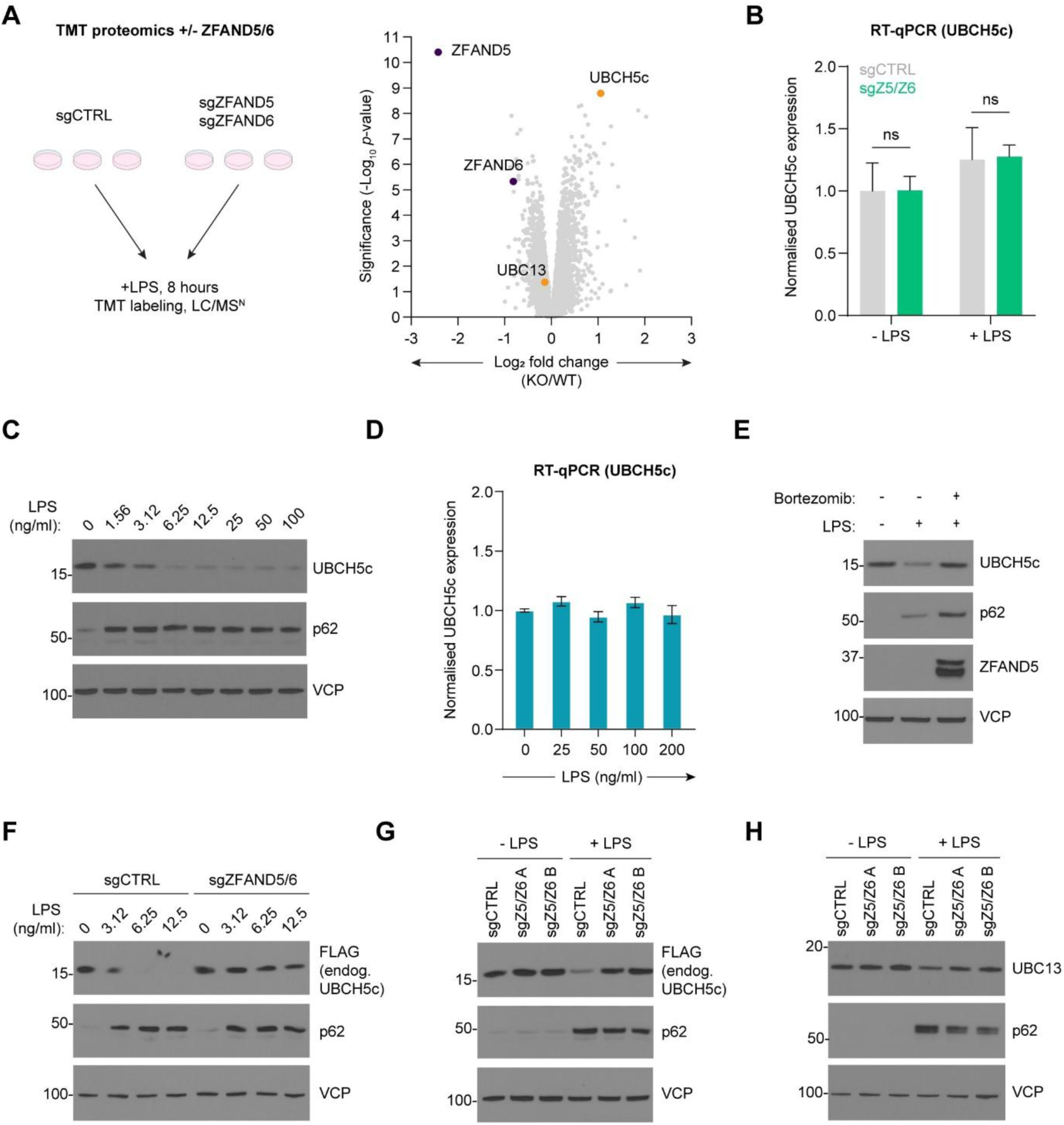
ZFAND5/6 regulate the abundance of UBCH5c. **(A)** Identification of candidate ZFAND5/6 substrates via quantitative proteomics. Control or ZFAND5/6 knockout cells (RAW264.7) were treated ± LPS (100 ng/ml) for 8 hours and analysed by quantitative proteomics. The ‘LPS treated’ samples are shown. KO: Knockout. WT: Wild-type. **(B)** UBCH5c is not transcriptionally upregulated in ZFAND5/6 knockout cells. RT-qPCR analysis of UBCH5c expression in cells cultured in parallel with the ones analysed in (A). Expression was normalised to GAPDH and presented as fold change relative to the untreated, non-targeting sgRNA control (sgCTRL). Error bars: mean ± SD. NS: Not significant (p > 0.05). **(C-E)** UBCH5c is degraded by the proteasome following LPS treatment. UBCH5c is downregulated in response to LPS treatment (C) and this effect is post-transcriptional, as indicated by RT-qPCR analysis of UBCH5c expression in similarly treated cells (D). Proteasome inhibition (Bortezomib, 100 nM) blocks the degradation of UBCH5c stimulated by LPS (10 ng/ml) (E). Cells were treated with LPS overnight. p62 and ZFAND5 were included as positive controls for LPS (Fujita *et al*., 2011) and Bortezomib treatment, respectively. In (D), expression was normalised to GAPDH and presented as fold change relative to the untreated control. Error bars: mean ± SD. **(F-H)** ZFAND5/6 regulate UBCH5c, but not UBC13. (F) ZFAND5/6 were deleted via CRISPR-Cas9 in cells expressing endogenously tagged UBCH5c-FLAG and the cells were treated with an LPS titration overnight. (G) As in (F) but with two independent sgRNA sets and a single LPS concentration (10 ng/ml). (H) As in (G) but the control E2 UBC13 was analysed instead of UBCH5c.

To determine whether UBCH5c is dynamically regulated during TLR4 signalling, we monitored its expression in LPS-treated cells. UBCH5c was markedly downregulated in response to LPS stimulation (**Fig. 5C**). The scaffold protein p62/SQSTM1 was used as a positive control for LPS treatment (Fujita *et al*, 2011) and will be investigated in a subsequent section. UBCH5c downregulation was post-transcriptional (**Fig. 5D**), dependent on the proteasome (**Fig. 5E**), and strongly impaired in ZFAND5/6 deficient cells (**Fig. S14B**). UBCH5c is the predominant member of the UBCH5a/b/c gene family, whose isoforms are highly similar but differentially regulated (Roman-Trufero & Dillon, 2022). Commercially available antibodies raised against UBCH5c may cross-react with UBCH5a and UBCH5b. To ensure that we were specifically analyzing UBCH5c, we used CRISPR-Cas9 to epitope tag the endogenous protein. ZFAND5/6 deficient cells showed an increase in the abundance of endogenous, FLAG-tagged UBCH5c at steady state and following LPS stimulation (**Fig. 5F-G**; **Fig. S14C**). In contrast, UBC13 was unaffected by loss of ZFAND5/6 (**Fig. 5H**). Finally, to determine whether ZFAND5/6 physically interact with UBCH5c, we used CRISPR-Cas9 to HA-tag endogenous ZFAND5 and immunoprecipitated the protein under non-denaturing conditions (**Fig. S14D**). UBCH5c co-immunoprecipitated with ZFAND5, indicating that the two proteins physically associate. Together, these results show that ZFAND5 and ZFAND6 regulate the abundance of UBCH5c, an abundant E2 enzyme with direct relevance to NF-κB signalling.

### ZFAND5 and p62 physically interact and convergently regulate the abundance of UBCH5c

A20/TNFAIP3 associates with UBCH5c via its A20^ZnF4^ domain; however, the domain is necessary but not sufficient to bind E2 enzymes (Bosanac *et al*., 2010; Martens & van Loo, 2020; Shembade *et al*., 2010). A20 forms a complex with an additional protein, TAX1BP1, which is required for E2 recruitment (Bosanac *et al*., 2010; Shembade *et al*., 2010). We therefore hypothesised that ZFAND5/6 might also form a complex with an additional protein that could stabilize or promote the interaction with UBCH5c. To identify candidate proteins, we immunoprecipitated endogenously tagged ZFAND5 from LPS-stimulated RAW264.7 cells and analysed the eluates by mass spectrometry. The joint top hit (as ranked by protein abundance) was p62/SQSTM1, a scaffold protein that regulates NF-κB signalling in multiple contexts and binds UBCH5c directly and specifically via a dedicated domain (**Fig. 6A**) (Duran *et al*, 2008; Peng *et al*, 2017; Sanchez-Martin *et al*, 2019; Sanz *et al*, 2000; Wooten *et al*, 2005; Zhong *et al*, 2016). The hits also included 19 proteasome subunits which served as internal positive controls and validated our approach.

**Figure 6.**
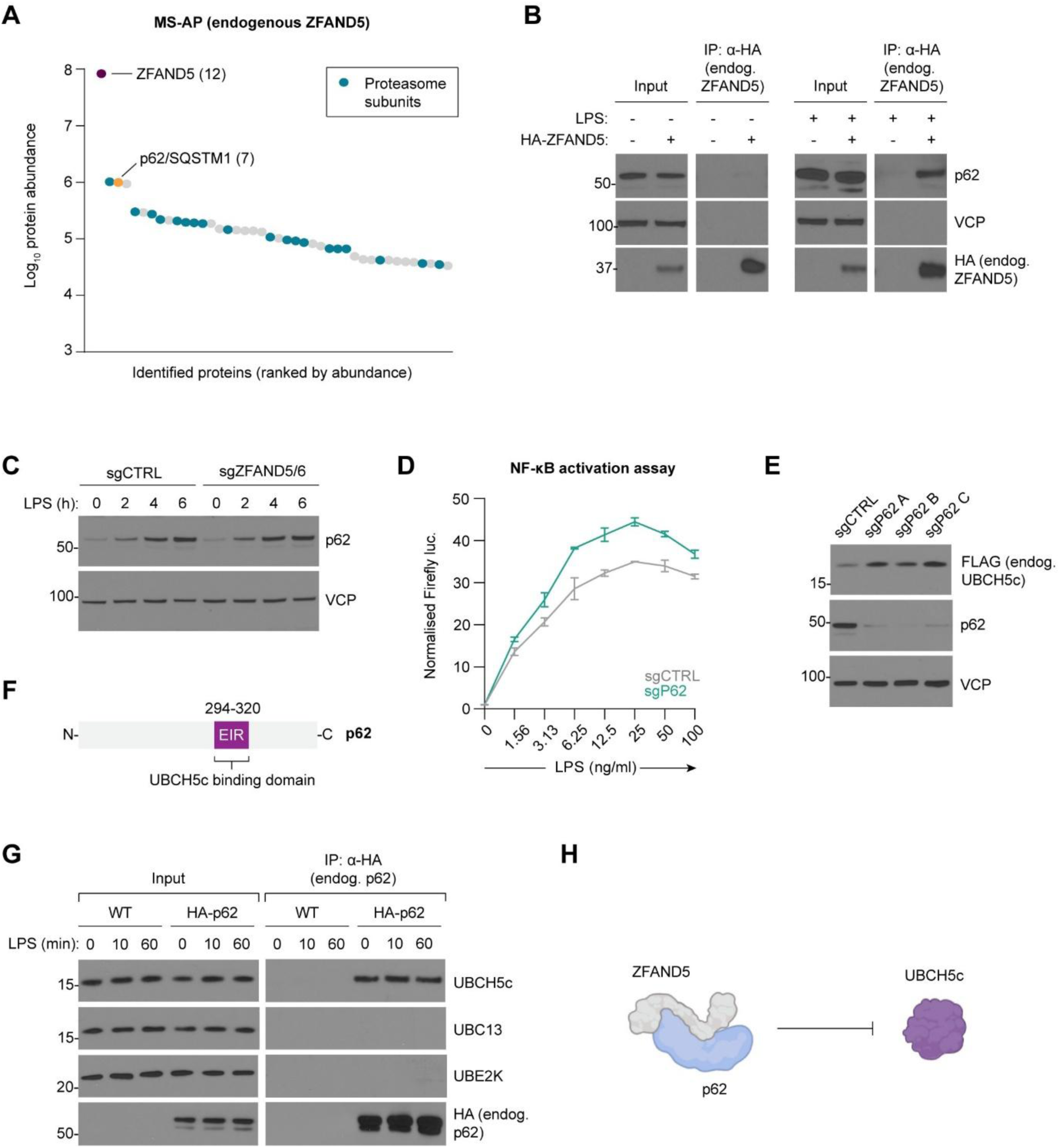
ZFAND5 and p62 physically interact and convergently regulate the abundance of UBCH5c. **(A-B)** ZFAND5 associates with p62 in an LPS-sensitive manner. (A) Identification of ZFAND5-binding proteins by MS-AP. Lysates from endogenously tagged HA-ZFAND5 and wild-type control cells (RAW264.7) treated with LPS overnight (10 ng/ml) were subjected to anti-HA immunoprecipitation and analysed by MS-AP. All proteins present in the control IP were excluded. Numbers to the right of the hits: unique peptides. (B) As in (A) but ± LPS as indicated (10 ng/ml, 17 hours) and the samples were resolved by immunoblotting. VCP: Loading and specificity control. **(C)** p62 is rapidly upregulated following LPS stimulation and it is not a ZFAND5/6 substrate. Control or ZFAND5/6 knockout cells were treated with LPS (10 ng/ml) as indicated. **(D)** p62 restrains NF-κB activation. p62 was knocked out in the NF-κB reporter cells and NF-κB activation was assayed across an LPS titration (6-hour treatment). Firefly signal was normalised against Renilla signal and presented as fold change relative to the untreated, non-targeting sgRNA control (sgCTRL). Error bars: mean ± SD from three technical replicates. **(E)** UBCH5c is upregulated in p62 knockout cells. p62 was deleted in cells expressing UBCH5c-FLAG (endogenously tagged). **(F)** Simplified p62 domain map highlighting the E2-interacting region (EIR). This domain directly and specifically binds to UBCH5c (Peng *et al*., 2017). **(G)** Validation of the p62/UBCH5c interaction. Cells expressing HA-p62 (endogenously tagged) or wild-type control cells (‘WT’) were treated with LPS as indicated (10 ng/ml), and subjected to anti-HA immunoprecipitation under non-denaturing conditions. UBC13 and UBE2K were included as specificity controls. **(H)** Model: ZFAND5 and p62 physically associate and convergently regulate UBCH5c.

We validated the ZFAND5/p62 interaction and found that it was strongly enhanced in LPS treated cells (**Fig. 6B**). This prompted us to ask whether p62 is, like ZFAND5, upregulated in response to LPS treatment. Consistent with a previous report (Fujita *et al*., 2011), p62 was rapidly upregulated at the protein level following LPS stimulation (**Fig. 6C**). In addition, p62 induction was unchanged in ZFAND5/6 knockout cells, suggesting that p62 is a cofactor and not a substrate of ZFAND5/6. Together, these results indicate that ZFAND5 and p62 physically associate in response to LPS stimulation. Next, we asked whether p62 regulates NF-κB signalling in the TLR4 pathway. Like ZFAND5/6, p62 deletion enhanced NF-κB activation in response to LPS stimulation (**Fig. 6D**). In addition, UBCH5c was constitutively upregulated in p62 knockout cells, indicating that ZFAND5/6 and p62 convergently regulate UBCH5c and may restrain NF-κB signalling through a similar mechanism (**Fig. 6E**). As p62 is a promiscuous scaffold protein, its function may be to promote or stabilize the interaction between ZFAND5/6 and UBCH5c. Consistent with this idea, a previous study found that p62 binds to UBCH5c directly and specifically via its dedicated E2-interacting region (EIR) (**Fig. 6F**) (Peng *et al*., 2017). The p62/UBCH5c interaction was readily confirmed by immunoprecipitation of endogenously tagged p62, which associated strongly and specifically with UBCH5c but not with control E2 enzymes UBC13 or UBE2K (**Fig. 6G**). Taken together, these results indicate that ZFAND5 and p62 physically interact and convergently regulate UBCH5c (**Fig. 6H**).

### ZFAND5/6 and p62 suppress TRAF6 activity

UBCH5c is recruited by several E3 ligases central to NF-κB signalling (Roman-Trufero & Dillon, 2022; Stewart *et al*, 2016). We therefore hypothesised that ZFAND5/6 and p62 might suppress the activation of one or more of these E3 ligases by degrading UBCH5c. TRAF6 was a strong candidate for several reasons: First, TRAF6 is essential for and central to NF-κB activation in the TLR4 pathway (Hu & Sun, 2016; Liu *et al*, 2017). Second, p62 physically associates with TRAF6 and modulates NF-κB signalling in the IL-1, NGF, and RANK pathways by modifying TRAF6’s activity (Duran *et al*, 2004; Sanchez-Martin *et al*., 2019; Sanz *et al*., 2000; Wooten *et al*., 2005). Third, A20 and TAX1BP1 extract UBCH5c (and UBC13) from TRAF6 to terminate NF-κB signalling in several pathways (Shembade *et al*., 2010). Finally, TRAF6 recruits UBCH5c and UBC13 to assemble its K63-linked polyubiquitin chains. This assembly requires UBCH5c to ‘prime’ chain synthesis by transferring the first ubiquitin to the substrate, while UBC13 mediates chain extension (Petroski *et al*, 2007; Roman-Trufero & Dillon, 2022; Shembade *et al*., 2010; Windheim *et al*, 2008). Priming is necessary because UBC13 (and other linkage-specific E2s) can extend but not initiate substrate-anchored ubiquitin chains (Stewart *et al*., 2016).

The interaction between p62 and TRAF6 is mediated by p62’s dedicated TRAF6-binding (TB) domain (**Fig. 7A**) (Sanz *et al*., 2000). Immunoprecipitation of endogenously tagged TRAF6 confirmed that TRAF6 and p62 physically interact in RAW264.7 cells (**Fig. 7B**). We also validated the interaction between TRAF6 and UBCH5c (**Fig. S15A**). The interaction was specific, as TRAF6 failed to associate with the control E2 enzymes UBE2Z, UBE2S, or UBE2B, and the TRAF6/UBCH5c interaction progressively dissociated following prolonged LPS stimulation.

**Figure 7.**
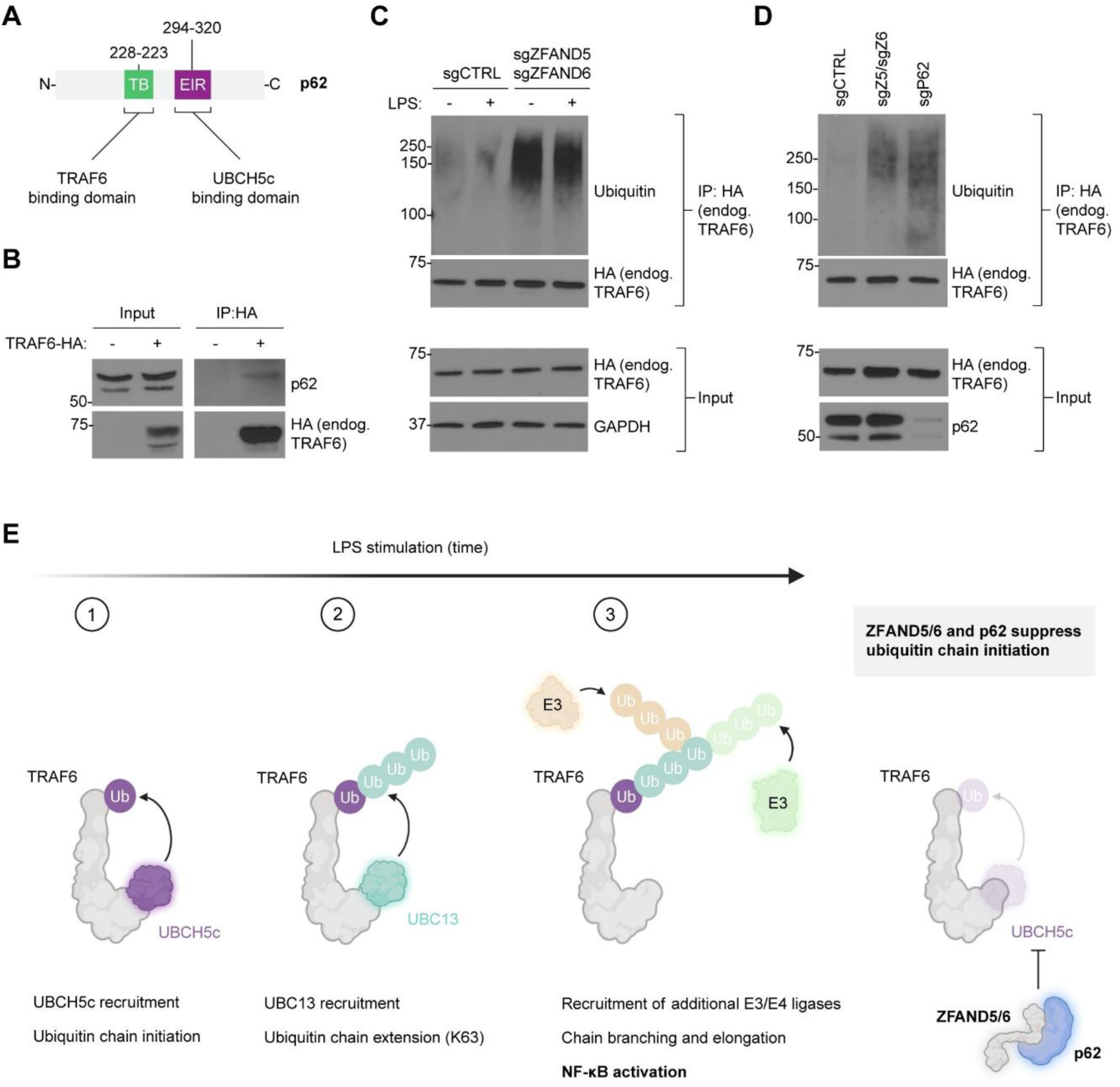
ZFAND5/6 and p62 suppress TRAF6 activity. **(A)** Domain map highlighting p62’s TRAF6 binding domain (TB) and E2-interacting region (EIR). **(B)** TRAF6 associates with p62. Wild-type cells, or cells expressing TRAF6-HA (endogenously tagged) were treated with LPS (10 ng/ml, 4 hours) and TRAF6-HA was immunoprecipitated under non-denaturing conditions. Symbols indicate wild-type (-) and knock-in (+). **(C-D)** TRAF6 activity is heightened in ZFAND5/6 and p62 knockout cells. Endogenously tagged TRAF6 was immunoprecipitated under denaturing conditions. Cells were treated (-/+) LPS for 7 minutes (C) or pretreated with LPS for 4 hours to boost p62 signal (D). 10 ng/ml LPS in both cases. **(E)** Model: ZFAND5/6 and p62 regulate TRAF6 activation and NF-κB signalling by regulating UBCH5c. During TLR4 signalling, TRAF6 recruits UBCH5c to prime ubiquitin chain assembly (step 1) before recruiting UBC13 multiple times to processively build K63-linked chains (step 2). Additional E3/E4 ligases extend and modify these chains to generate large, branched ubiquitin assemblies containing multiple linkage types which drive NF-κB activation (step 3). ZFAND5/6 and p62 suppress TRAF6’s activity by regulating UBCH5c, thus suppressing the priming of ubiquitin chain synthesis (step 1).

To determine whether ZFAND5/6 suppress TRAF6’s activity, we immunoprecipitated endogenously tagged TRAF6 from control and ZFAND5/6-deficient cells under denaturing conditions. TRAF6-associated ubiquitination was markedly increased in ZFAND5/6 knockout cells both at steady state and following LPS stimulation (**Fig. 7C**; **Fig. S15B**). Similar results were observed in p62 knockout cells (**Fig. 7D**). These findings, together with the observed interactions between ZFAND5, p62, UBCH5c, and TRAF6, suggest a model whereby ZFAND5/6 and p62 restrain TRAF6 activity, and thus NF-κB activation, by regulating UBCH5c (**Fig. 7E**).

## Discussion

It has long been appreciated that some proteins can be targeted to the proteasome without ubiquitination, but the prevalence of this phenomenon is unclear, and the underlying mechanisms are poorly understood. Here, we used pulsed-SILAC proteomics to identify both known and novel *UbInPD* proteins. We identified a significant number of putative *UbInPD* proteins (*n* = 231), suggesting that *UbInPD* may make a greater contribution to proteostasis than has previously been recognised. Similar conclusions were reached by Makaros and colleagues, who recently presented two screens for *UbInPD* proteins using a genetic reporter system (Makaros *et al*., 2023). Our results complement and extend this work by systematically identifying *UbInPD* proteins in an endogenous cellular context.

Among the highest confidence hits were the proteasome activation/shuttle factors ZFAND5 and ZFAND6. We showed that ZFAND5 is rapidly degraded by an intrinsic, ubiquitin-independent degron that can allosterically activate the proteasome as it engages (Lee *et al*., 2018; Lee *et al*., 2023). These observations reveal a direct coupling between proteasome activation and ZFAND5 degradation and support a model in which ZFAND5 (and presumably ZFAND6) function within a feedback circuit that limits excessive proteasome activation. We also note conceptual similarities between ZFAND5 and the yeast transcription factor Rpn4, which promotes the production of new proteasomes and is also rapidly degraded by an intrinsic, ubiquitin-independent degron (Ha *et al*., 2012; Xie & Varshavsky, 2001).

Beyond proteostasis, our findings identify ZFAND5/6 as negative regulators of inflammatory signalling. We found that ZFAND5/6 restrain NF-κB activation in the TLR4 pathway and regulate the abundance of the E2 ubiquitin-conjugating enzyme UBCH5c. This was notable because UBCH5c functions as a promiscuous ‘priming’ E2 for multiple E3 ligases, including TRAF6, and has established roles in NF-κB signalling (Roman-Trufero & Dillon, 2022; Shembade *et al*., 2010; Windheim *et al*., 2008; Xu *et al*, 2009). The selective regulation of UBCH5c, but not the chain-extending E2 UBC13, is consistent with a model in which ZFAND5/6 influence inflammatory signalling through the control of ubiquitin chain initiation rather than elongation.

Mechanistically, our data suggest that ZFAND5/6 function together with the scaffold protein p62. ZFAND5 associates with p62 in an LPS-dependent manner, and both proteins regulate UBCH5c abundance and NF-κB signalling. This parallels the organisation of the A20/TAX1BP1 complex, which also targets E2 enzymes to suppress NF-κB activation (Shembade *et al*., 2010). As p62 binds to UBCH5c directly (Peng *et al*., 2017), its role in this context may be to promote or stabilise the interaction between ZFAND5/6 and UBCH5c. However, the precise architecture of this complex remains to be defined.

Consistent with a role in regulating upstream signalling, TRAF6 activity is enhanced in ZFAND5/6- and p62-deficient cells. Given that TRAF6 depends on UBCH5c to initiate ubiquitin chain assembly, these findings support a model in which ZFAND5/6 and p62 restrain NF-κB activation by limiting UBCH5c availability. Whether ZFAND5/6 and p62 preferentially target UBCH5c when it is in complex with TRAF6 remains to be determined, but is suggested by the direct interaction between p62 and TRAF6 and by the precedent set by A20 and TAX1BP1 (Shembade *et al*., 2010).

Taken together, our findings expand the known landscape of ubiquitin-independent proteasomal degradation, provide new insights into the regulation of proteasome activation and proteostasis, and reveal unexpected links between proteasome targeting, ubiquitin chain priming, and inflammatory signalling.

## Acknowledgments

We thank Professor Clare Bryant (University of Cambridge) for gifting the iBMDM cells, and Professor Felix Randow (University of Cambridge) for his critical reading of the manuscript.

SJW, CM, JCW, CMG, AD, and PJL were funded by a Wellcome Trust Discovery Award (227418/Z/23/Z) and a grant from CRUK (22492).

PG and NM are supported by the Evelyn Trust (24/55 (Med-24-2316)).

GLG, NW, and JAN were funded by the Wellcome Trust (215477/Z/19/Z).

RTT is a Pemberton-Trinity Fellow supported by an Academy of Medical Sciences Springboard Grant (SBF007\100019), an Isaac Newton Trust/Wellcome ISSF/University of Cambridge Joint Research Grant and an ERC Starting Grant (ERC-2024-STG 101160971).

We also acknowledge the NIHR Cambridge Biomedical Research Centre (BRC).

## Author contributions

Conceptualisation: SJW, PJL, RTT, NJM

Methodology: SJW, JCW, PG, GLG, CM, AD, CMG, NW

Investigation: SJW, JCW, PG

Funding acquisition: PJL, JAN, NJM

Supervision: PJL

Writing: SJW, PJL, JAN

## Methods

### Cell culture

HeLa (RRID:CVCL_0030), HEK293T (RRID:CVCL_0063), RAW264.7 (RRID:CVCL_C6XG), and immortalised bone marrow-derived macrophages (iBMDMs) were cultured in Dulbecco’s Modified Eagle’s Medium (DMEM) supplemented with 10% fetal calf serum (FCS) and penicillin/streptomycin. The iBMDM cells were a kind gift from Professor Clare Bryant (University of Cambridge, UK). All cells were cultured at 37°C with 5% CO_2_. The HeLa, HEK293T and ‘wild-type’ RAW264.7 cells were sourced from ATCC and the RAW264.7 NF-κB reporter cell line was sourced from BPS Bioscience (#79978). The reporter cell line was further engineered to express constitutive Renilla luciferase as will be described in a subsequent section. All cell lines were tested for mycoplasma every month (Lonza, LT07-118).

### Chemicals

Cells were treated with TAK-243 (Selleckchem, S8341, 4.5 µM), Bortezomib (Selleckchem, S1013; 100 nM), and cycloheximide (Cell Signaling Technology, #2112; 10 µg/ml) where indicated. Cells were treated with ultrapure LPS as indicated (InvivoGen, tlrl-pb5lps). The ultrapure LPS exclusively stimulates the TLR4 pathway.

### RT-qPCR

RNA was extracted from approximately 1 × 10^6^ cells via the RNeasy Plus Kit (Qiagen, #74134). 0.5 to 1 µg RNA was used as template for cDNA synthesis. SuperScript III (Invitrogen, #18080093) was used according to the manufacturer’s instructions, and OligoDT(15) (Promega, C110A) was used to selectively amplify the polyadenylated (mRNA) fraction of the samples. The cDNA samples were diluted 1:5 in nuclease-free water and the target transcripts were amplified using SYBR Green PCR Master Mix (ThermoFisher, #4309155). Samples were run in technical triplicate using a PCR thermocycler (Applied Biosystems, 7500 Real Time PCR System). The ΔΔCT method was used to calculate relative transcript abundance, and GAPDH was used as an internal control. Primer details are shown below.

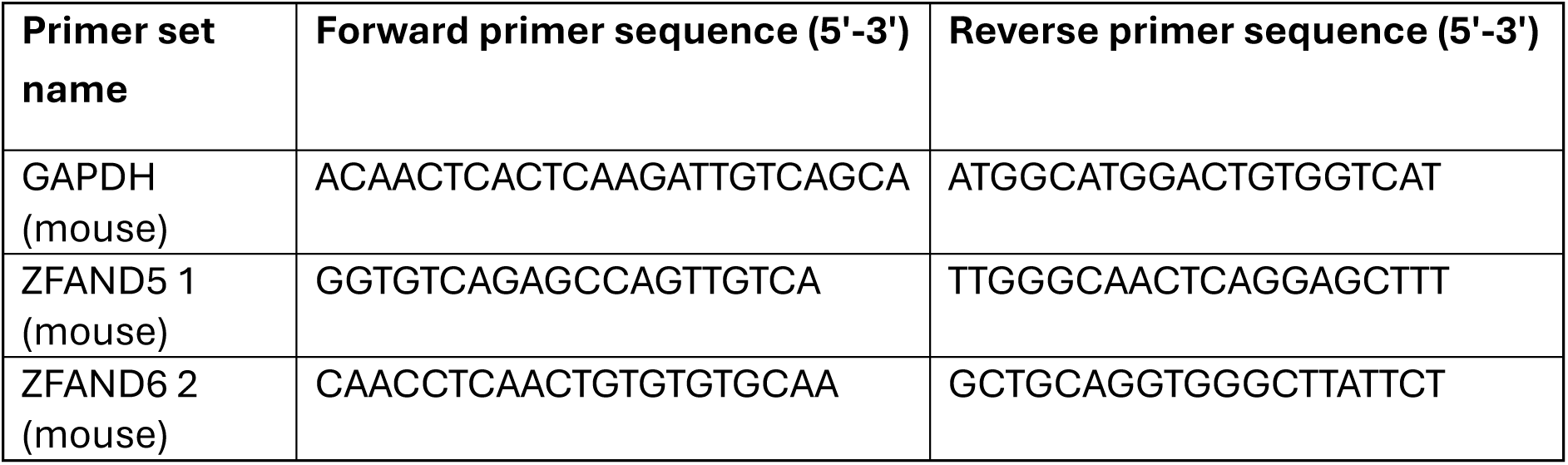
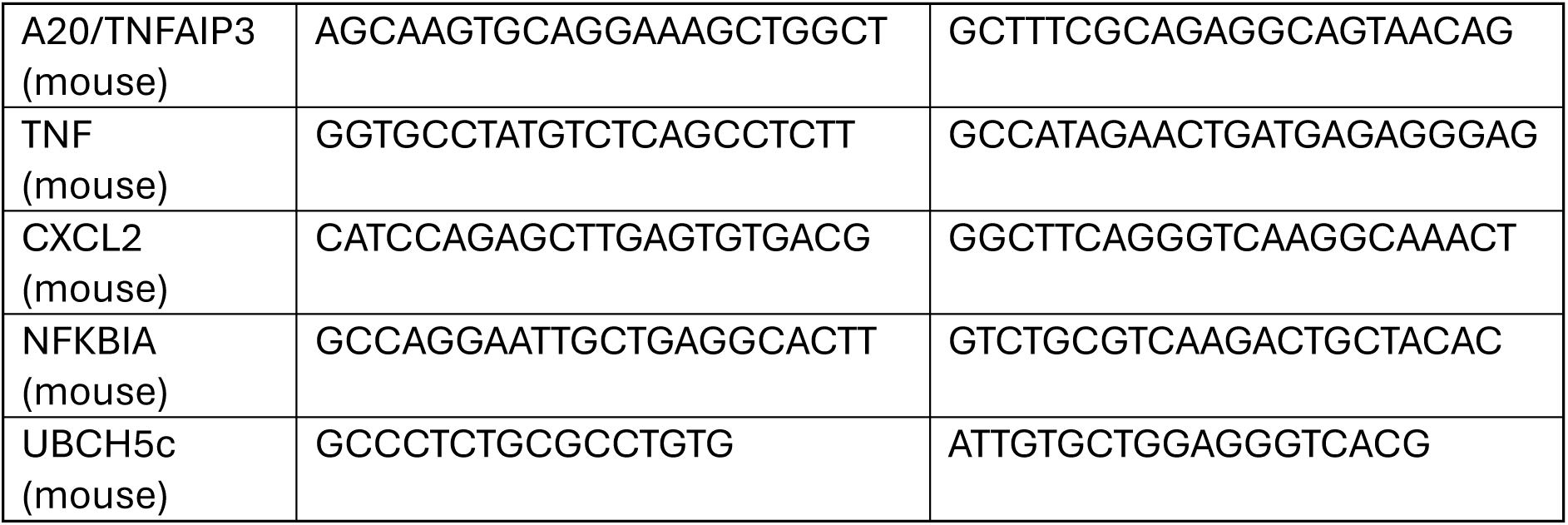

### Plasmids

For the UBA6 knockout experiment, single guide RNAs were cloned into pSpCas9(BB)−2A-Puro V2 (Addgene #62988) using the BpiI restriction enzyme (Ran *et al*, 2013). This is an ‘all-in-one’ plasmid that encodes both SpCas9 and an sgRNA scaffold. Primers used in the cloning procedures are shown in the table below. For the various lentiviral expression plasmids, gBlocks (double-stranded DNA fragments, IDT) or PCR products were restriction cloned into pHRSIN-P_SFFV_-P_PGK_-Hygromycin^R^ or pHRSIN-P_SFFV_-P_PGK_-Puromycin^R^, or custom variants of these plasmids that append an N-terminal 3xHA tag to the encoded insert (Demaison *et al*, 2002). BamHI/KpnI/NotI restriction enzymes were used. ZFAND5, ZFAND6, and the ΔAN1 and M1 mutants were ordered as gBlocks, whereas the ΔA20 mutants and AN1/CTR fragments were amplified from the cloned gBlocks using the primers shown below. The ZFAND5 A20 ‘M1’ point mutant (C30A and C33A) has been described previously (Hishiya *et al*, 2006; Lee *et al*., 2018). The truncation mutants lacked the A20 or AN1 domains as defined: ZFAND5 A20 (residues 8-42), ZFAND5 AN1 (residues 148-194), ZFAND6 A20 (residues 8-42), ZFAND6 AN1 (residues 143-189). For the GPS plasmids, ODC1, AUH, ZFAND5, ZFAND6, and the ZFAND5/6 fragments were ordered as gBlocks with 5’ and 3’ Gateway overhangs, and cloned into pHAGE-GPS3.0-DEST (Yen *et al*, 2008) via the Gateway system (Thermo Fisher Scientific). Renilla luciferase was amplified from pGloSensor-30F (Promega) as previously described (Gerber *et al*, 2022).

#### Primers used to clone sgRNA constructs

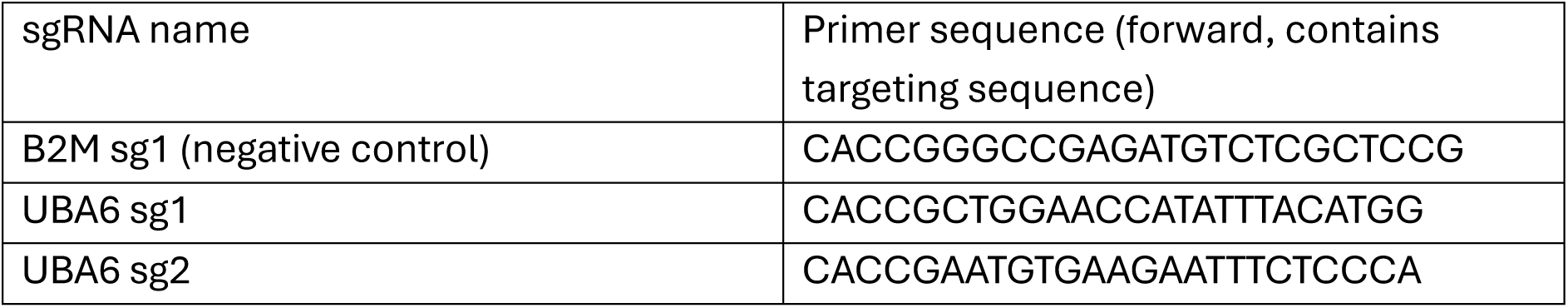

#### Primers used to clone lentiviral expression constructs

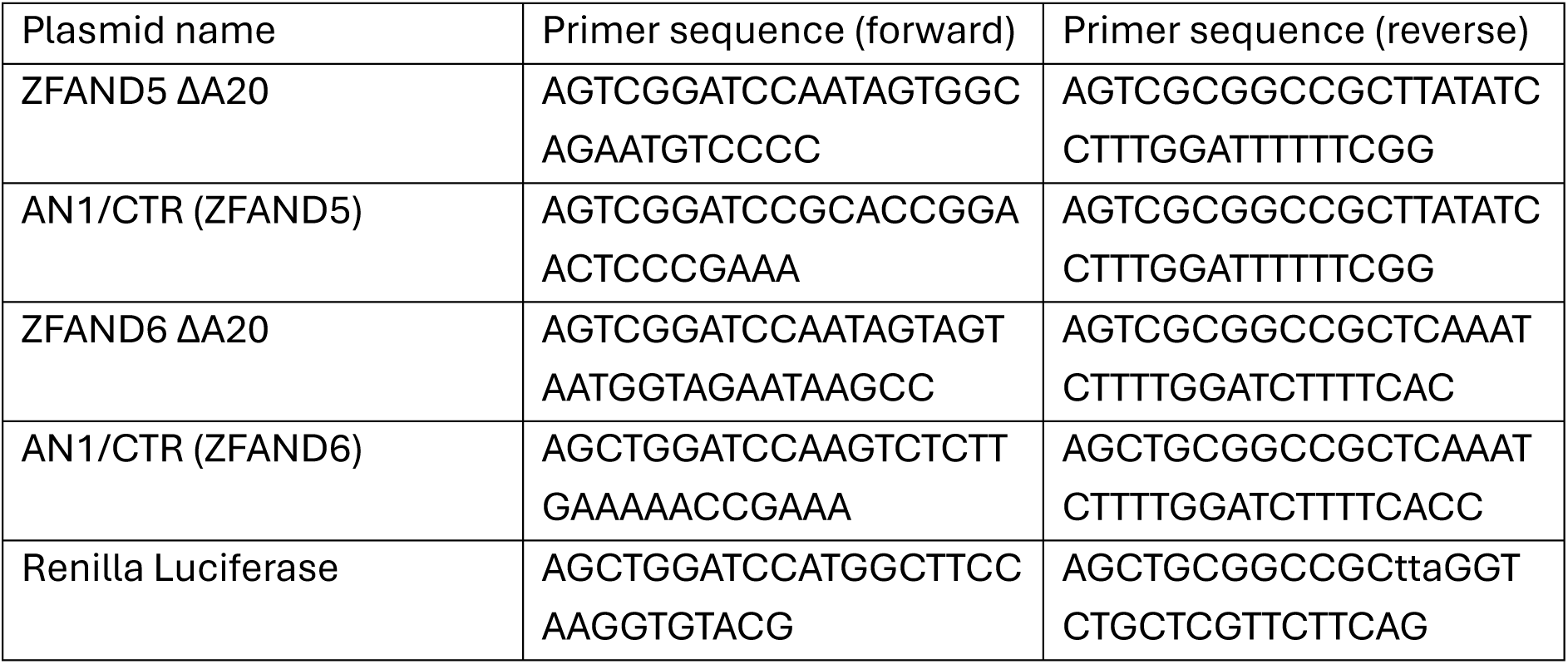

### Antibodies

| Antibody | Supplier | Catalogue ID | RRID | Working concentration |
| --- | --- | --- | --- | --- |
| α-VCP | Abcam | ab109240 | RRID:AB_10862588 | 1:10,000 |
| α-Ubiquitin | Life Sensors | VU101 | RRID:AB_2716558 | 1:1000 |
| α-ADIPOR2 | ProteinTech | 14361-1-AP | RRID:AB_2222052 | 1:1000 |
| α-HMGCR | Santa Cruz | sc-271595 | RRID:AB_10650274 | 1:1000 |
| α-HA | Cell Signalling Technology | 2999 | RRID:AB_1264166 | 1:5000 |
| α-HA | Roche | 11867423001 | RRID:AB_390918 | N/A (IP only) |
| α-ZFAND5 | Abcam | ab272580 | - | 1:1000 |
| α-ZFAND6 | Novus | Nbp1-90060 | RRID:AB_11003101 | 1:1000 |
| α-UBCH5c | Proteintech | 11677-1 | RRID:AB_2256691 | 1:1000 –<br>1:100,000 |
| α-p62 | Cell Signalling Technology | 5114 | RRID:AB_10624872 | 1:5000 |
| α-A20 | Cell Signalling Technology | 5630 | RRID:AB_10698880 | 1:1000 |
| α-GAPDH | Cell Signalling Technology | 2118 | RRID:AB_561053 | 1:5000 |
| α-UBC13 | Cell Signalling Technology | 6999 | RRID:AB_10828936 | 1:10,000 |
| α-UBE2K | Proteintech | 11834-3-AP | RRID:AB_2241168 | 1:1000 |
| α-UBE2Z | Proteintech | 16928-1-AP | RRID:AB_2211782 | 1:1000 |
| α-UBE2B | Proteintech | 10733-1-AP | RRID:AB_2212174 | 1:1000 |
| α-UBE2S | Proteintech | 14115-1-AP | RRID:AB_2256809 | 1:4000 |
| Direct-Blot™ HRP<br>anti-<br>DYKDDDDK<br>Tag (FLAG) | BioLegend | 637312 | RRID:AB_2734514 | 1:10,000 |
| TrueBlot™<br>Anti-Rabbit-HRP | Jackson<br>ImmunoResearch | 18-8816-31 | RRID:AB_2610847 | 1:10,000 |
| TrueBlot™<br>Anti-Mouse-HRP ULTRA | Jackson<br>ImmunoResearch | 18-8817-30 | RRID: AB_2610849 | 1:10,000 |

### CRISPR/Cas9 (CRISPRko)

For the UBA6 knockout, HeLa cells were transfected with the UBA6-targeting plasmids or a non-targeting control. The TransIT-HeLaMONSTER transfection kit was used according to the manufacturer’s protocol (Mirus). The targeting sequences are shown in the ‘plasmids’ section.

For the RAW264.7 knockouts, Alt-R® CRISPR-Cas9 reagents were ordered (IDT) and delivered via nucleofection (Lonza, 4D-Nucleofector system). Predesigned crRNAs or sgRNAs targeting murine ZFAND5, ZFAND6, p62, and A20, were ordered from IDT, along with non-targeting control crRNAs and sgRNAs. When crRNAs were ordered, crRNAs and tracrRNAs were annealed before assembly of the ribonucleoprotein (RNP) complexes. Equimolar (200 pmol) mixtures of crRNA and tracrRNA (IDT, #1073191) were suspended in Duplex Buffer (IDT, #1072570) to a final volume of 4.5 µl, heated to 95°C for 5 minutes, then allowed to anneal at room temperature for at least 40 minutes. To assemble the RNP complexes, 5 µg of purified Cas9 protein (IDT, #1081059) and 2.2 µl electroporation enhancer (IDT, #1075916) were added to the sgRNAs and the samples were incubated at room temperature for 15 minutes before being cooled on ice.

RAW264.7 cells were harvested by scraping, and 1 × 10⁶ cells were nucleofected with RNP complex in each reaction. The SF Cell Line kit (Lonza, V4XC-2024) was used according to the manufacturer’s instructions and the preprogrammed RAW264.7 nucleofection protocol was used. Knockout efficiency was confirmed by immunoblotting. The knockout efficiency was such that populations, not clones, were analyzed in all cases. The crRNA/sgRNA sequences are shown below.

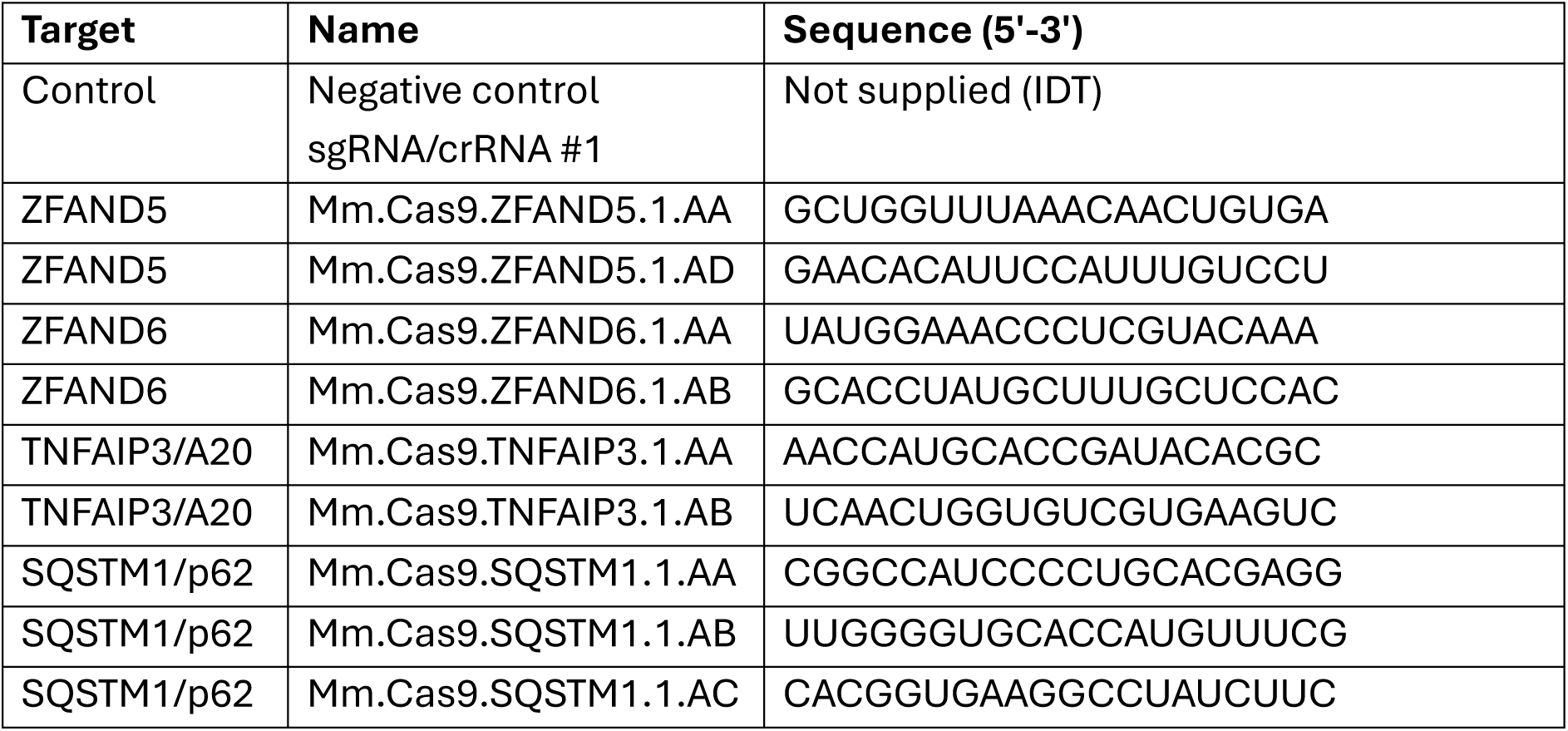

### CRISPR/Cas9 (CRISPR knock-ins)

For the CRISPR-Cas9-mediated knock-ins, the Alt-R® CRISPR-Cas9 system (IDT) was used with certain modifications. The following RAW264.7 knock-in cell lines were generated: HA-ZFAND5, UBE2D3-FLAG (UBCH5c), HA-SQSTM1 (p62), and TRAF6-HA. Predesigned crRNAs or sgRNAs targeting the 5’ or 3’ ends of the genes’ coding sequences were ordered from IDT, as were custom ssODN repair templates that incorporated the indicated tags. RNP complexes were assembled as described in the previous section, and 0.5 nmol ssODN repair template was added to the complexes immediately prior to nucleofection. After nucleofection, cells were transferred to a 6-well plate and cultured in DMEM containing AZD-7648 (1 µM), a DNA-PK inhibitor (Selleckchem, S8843). Cells were cultured at 32°C for 24 hours, then transferred to regular DMEM and cultured at 37°C as before. Knock-ins were validated by gDNA PCR, Sanger sequencing, and immunoblotting. For the HA-ZFAND5, HA-SQSTM1 (p62), and TRAF6-HA cell lines, clones were isolated by serial dilution. For the UBE2D3-FLAG (UBCH5c) cell line, knock-in efficiency was so high that the bulk nucleofected population was taken forward for further use. The crRNA/sgRNA sequences and ssODN repair template sequences are shown below.

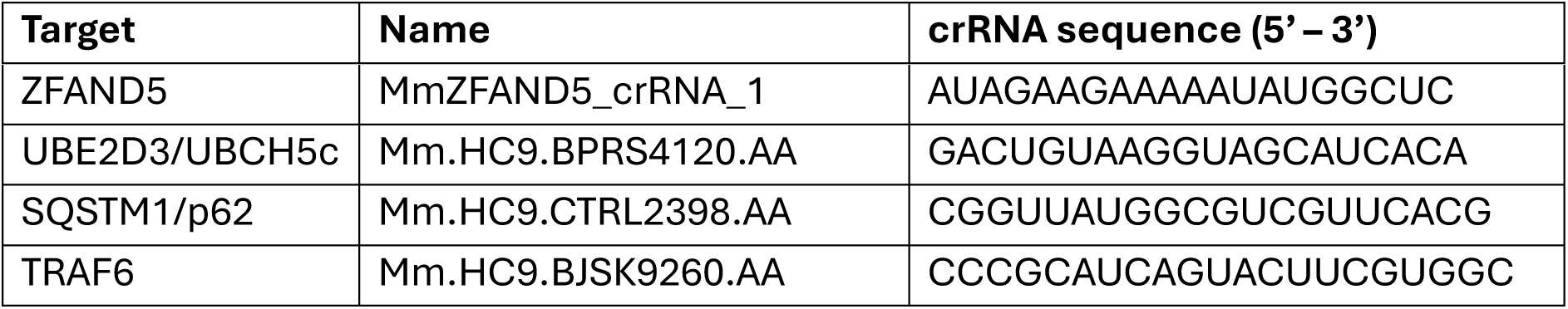

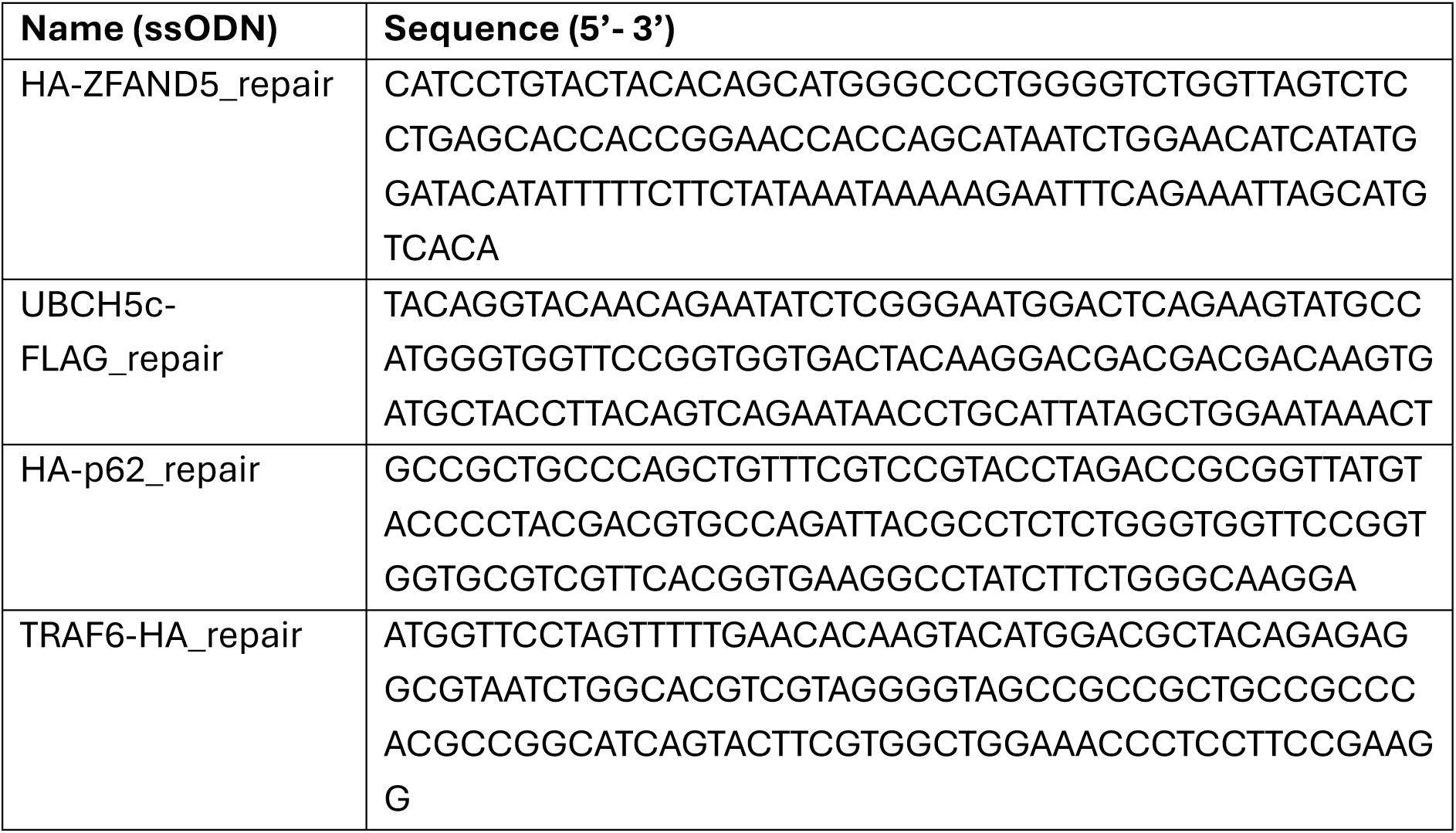

### Luciferase assays

For the NF-κB signaling assays, a commercially available RAW264.7 reporter cell line was obtained (BPS Bioscience, #79978) and further engineered as described in the ‘Lentiviral production and transduction’ section. This cell line stably expresses Firefly luciferase under the control of an NF-κB responsive promoter. The promoter consists of a minimal promoter and four tandem NF-κB response elements (5’-GGAATTTCC-3’). We further edited this cell line to stably express Renilla luciferase under the control of a strong, constitutive promoter (SFFV). The Renilla signal is a control for subtle differences in cell seeding density and lysis efficiency between the samples. We then used CRISPR/Cas9 to knock out ZFAND5/6 or other genes in the cell line. Knockout efficiency was verified by immunoblotting in every case. In all cases, populations, not clones, were analyzed.

After validation of knockout efficiency, the cells were assayed for NF-κB activation. 96-well plates were seeded with the various cell lines at an initial seeding density of approximately 30,000 cells/well. The next day, the cells were treated with an LPS titration for 5-6 hours, or with 10 ng/ml LPS across a 6-hour time-course. Each cell line/treatment combination was assayed in technical triplicate. At the end of the experiment, the media was removed from each well and replaced with Glo Lysis Buffer (Promega, E2661), and the plates were incubated at room temperature for 10 minutes with gentle agitation. The lysates were transferred to white 96-well plates and Firefly and Renilla luciferase signal was assayed sequentially using the Dual-Glo kit (Promega, E2920) and a luminometer. Each Firefly reading was normalized against the corresponding Renilla reading, and the resulting values were all normalized against the first sgCTRL, 0 ng/ml LPS sample.

### RNA-seq experiment

For the RNA-seq experiment, ZFAND5/6 were deleted in RAW264.7 cells via nucleofection. Populations, not clones, were analysed. Five days post-nucleofection, the control and ZFAND5/6-deficient cells were treated with LPS (1 ng/ml) across an 8-hour time-course. Samples were harvested and RNA was extracted using a Monarch Total RNA Miniprep kit (New England Biolabs, #T2010S). For the data analysis, paired-end raw sequencing data (FASTQ format) was first adapter trimmed using Trim Galore (https://doi.org/10.5281/zenodo.7598955) with standard settings. Salmon (Patro *et al*, 2017) was used to quantify transcripts from the trimmed data against the mouse genome (Gencode M38, GRCm39 (Mudge *et al*, 2025)) using a decoy-aware transcriptome index for accurate mapping. A mapping rate of >88% was observed for all samples. R was used to create a normalised count table from the Salmon transcript quantification files: data were loaded with tximport (Soneson *et al*, 2015) and counts were normalised using DESeq2 (Love *et al*, 2014). The RNA-seq dataset has been deposited in the Gene Expression Omnibus (GEO) under accession number GSE334096.

### Lentiviral production and transduction

To make the viruses, 600,000 HEK-293T cells were reverse transfected with three plasmids: the pHRSIN plasmid containing the gene of interest and two packaging plasmids (pCMVDR8.91 and pMD.G). The plasmid ratios were 1.0:0.65:0.35 respectively and the Trans-IT 293 kit was used (Mirus, MIR 2706). The supernatant was harvested three days post-transfection and sterile filtered (0.45 µm).

Cells (HeLa and RAW264.7) were transduced as follows: A 12-well plate was seeded with 200,000 cells/well and two-fold serial dilutions of filtered virus were added to each well. After three days, the cells were transferred to a 6-well plate and selected with Hygromycin or Puromycin for a further two to three days. Surviving cells treated with a relatively low volume of virus (as judged by the extent of cell death during selection) were taken forward for further analysis. For the RAW264.7 reporter cells, the ‘medium multiplicity of infection’ cell line was taken forward, as described in the main text. Renilla expression was confirmed by luciferase assay with the Dual-Glo kit (Promega, E2920). Renilla luciferase signal was robust and did not noticeably alter the Firefly luciferase signal.

### Flow cytometry analysis

HeLa cells stably overexpressing the GPS plasmids were seeded on 6-well plates to an initial density of 500,000 cells/well. The dual colour GPS plasmid reports on protein stability. After 24 hours, the cells were treated with DMSO, TAK-243 (4.5 µM) or Bortezomib (100 nM). After 17 hours, the cells were harvested via trypsinization, washed with ice-cold PBS, and resuspended in ice-cold PBS supplemented with 10% (v/v) FCS. Cells were phenotyped via an LSR Fortessa (BD) flow cytometer. At least 10,000 live cells were phenotyped for each sample and the data were analysed in FlowJo. The DsRed signal reports on transcription and translation, and the GFP signal reports on the abundance of the fusion protein. The GFP/DsRed ratio (which is a direct measure of protein stability) was calculated per cell and the results were normalised to the modal value and plotted as histograms. The same flow cytometer settings were used to collect data on each sample.

### Immunoprecipitation

For the MS-AP experiment, RAW264.7 wild-type and HA-ZFAND5 knock-in cells were seeded (two 15 cm dishes per condition) and treated with LPS overnight (10 ng/ml) followed by further treatment with LPS (10 ng/ml) and Bortezomib (100 nM) on the day of the experiment (4 h). Cells were lysed in Lysis Buffer (Tris-HCl, pH 7.4, 150 mM NaCl, 1% (v/v) Triton X-100, 1.5 mM MgCl_2_, 100 nM Bortezomib, 1x Protease Inhibitor Cocktail (Roche #11836170001)) and clarified by centrifugation. Lysates were normalized by BCA assay and precleared with 50 µl Protein G magnetic beads (Dynabeads Protein G, ThermoFisher, #10004D). HA-ZFAND5 was immunoprecipitated using 50 µl Protein G magnetic beads preconjugated with anti-HA antibody (Roche, #11867423001, 2 µg/sample). All preclearance, immunoprecipitation, and wash steps were carried out at 4°C with continuous overhead rotation. Beads were washed five times in Wash Buffer (Tris-HCl, pH 7.4, 1.5 mM MgCl_2_, 10% (v/v) Glycerol, 0.1% (v/v) NP-40, 100 nM Bortezomib, 1x Protease Inhibitor Cocktail (Roche #11836170001)) and bound proteins were eluted in Elution Buffer (Tris-HCl, pH 7.4, 1% (w/v) SDS). For the co-IP experiment validating the association between ZFAND5 and p62, the same conditions were used, except that an ‘LPS negative’ co-IP was carried out in parallel. These samples were treated with Bortezomib (100 nM) for 4 hours on the day of the IP but not with LPS at any time. For all other co-IP experiments, similar conditions were used, except that directly conjugated anti-HA beads were used (ThermoFisher, #88837) alongside similar Protein G beads used in the preclearance steps (ThermoFisher, #88848).

For the TRAF6 activity assays, endogenously tagged TRAF6-HA was immunoprecipitated under denaturing conditions. Cells were seeded in 6-well format and treated with LPS (10 ng/ml) as indicated. Samples were harvested on ice, lysed in denaturing Lysis Buffer (Tris-HCl, pH 7.4, 150 mM NaCl, 0.25% (w/v) Deoxycholic acid, 1% (v/v) NP-40, 1 mM EDTA, 1% (w/v) SDS, 0.5 mM PMSF, 2 mM MgCl_2_, 1x Protease Inhibitor Cocktail (Roche #11836170001)) and boiled (10 minutes, 95°C). The samples were cooled, Benzonase treated (Sigma, E1014), and diluted 1:10 to reduce the SDS concentration. Protein concentrations were normalised via BCA assay, and lysates were precleared with 10-30 µl Protein G beads (ThermoFisher, #88848). TRAF6-HA was immunoprecipitated using anti-HA magnetic beads (ThermoFisher, #88837). Samples were then washed 5-6 times in stringent Wash Buffer (Tris-HCl, pH 7.4, 500 mM NaCl, 0.25% (w/v) Deoxycholic acid, 1% (v/v) NP-40, 1 mM EDTA, 1x Protease Inhibitor Cocktail (Roche #11836170001)) and bound proteins were eluted in Elution Buffer (Tris-HCl, pH 7.4, 1% (w/v) SDS).

### Immunoblotting

Cells were collected by scraping into cold PBS, pelleted (600 × g, 4 minutes) and lysed in 1x RIPA buffer (50 mM Tris-HCl, pH 7.4, 150 mM NaCl, 0.25% (w/v) Deoxycholic acid, 1% (v/v) NP-40, 1 mM EDTA) supplemented with 1-5% (w/v) SDS, protease inhibitor cocktail (Roche #11836170001), and Benzonase (Sigma, E1014). The samples were incubated at room temperature for 5-15 minutes (for the benzonase), then heated at 55°C for 15 minutes. Typically, the concentration of total protein was determined for each sample using a BCA assay kit (ThermoFisher, #23227). Samples were normalized to a standard protein concentration, supplemented with concentrated Laemmli buffer (+DTT) and boiled (5-10 minutes, 80-95°C).

Protein samples were separated by SDS-PAGE using hand-cast, single percentage (8 to 12%) polyacrylamide gels, or on precast gradient (4-12%) Bis-Tris gels (ThermoFisher, NP0321; NP0322). For hand-cast gels, TGS running buffer was used (Tris-HCl, pH 8.3, 192 mM glycine, 0.1% (w/v) SDS). For precast gels, MOPS (ThermoFisher, #NP000102) or MES (ThermoFisher, #NP0002) running buffers were used to resolve high and low molecular weight proteins, respectively. Proteins were transferred to PVDF membranes (Merck, IPVH00010) that had been pretreated with methanol. For hand-cast gels, TGS transfer buffer was used (Tris-HCl, pH 8.3, 192 mM glycine, 0.1% (w/v) SDS, 20% (v/v) methanol). For precast gels, 1x NuPAGE transfer buffer supplemented with 10-20% (v/v) methanol was used (ThermoFisher, NP00061). In all cases, the membranes were blocked with 5% milk + PBS-T (PBS supplemented with 0.2% (v/v) Tween-20) for 30 minutes. The membranes were incubated with primary antibodies suspended in blocking buffer overnight at 4°C with continuous movement. The following day, the membranes were washed with PBS-T three times (5 minutes, continuous movement), then typically incubated with secondary antibodies suspended in milk for at least 1.5 hours (room temperature, continuous movement). The membranes were washed three times (5 minutes, continuous movement) and protein bands were visualized using ECL detection reagents (ThermoFisher, #32106; #34580; #34076; #34095). Bands were quantified using ImageJ.

### Quantitative proteomics

All chemicals were of AR grade or better and purchased from Merck/Sigma unless otherwise stated.

#### SILAC labelling

The SILAC labelling was carried out as previously described (Greenwood *et al*, 2016). HeLa cells were passaged for at least 7 divisions in either ‘heavy’ or ‘medium’ SILAC media. The media was SILAC DMEM (Thermo Fisher Scientific) supplemented with dialysed FBS (Thermo Fisher Scientific), penicillin/streptomycin, proline (Sigma, 280 mg/L) and either ‘heavy’ or ‘medium’ isotopically labelled lysine and arginine (Cambridge Isotope Laboratories, 50 mg/L). At the start of the experiment, the media was replaced with the same media supplemented with ‘light’ labelled lysine and arginine (Sigma, 50 mg/L) and either Bortezomib (Selleckchem) or TAK-243 (Selleckchem). The cells were harvested after 17 hours of drug treatment.

#### Sample preparation

For the ZFAND5/6 KO experiments, excess PBS was removed from cell pellets before resuspending the cells in 71 µL Resuspension Buffer (76 mM TEAB pH 8.5, 3 mM MgCl_2_, Benzonase (1400u/mL), 7 mM TCEP, 28 mM chloroacetamide) by pipetting. 25 µL 20% (w/v) SDS was then immediately added to the cell suspension using a low retention pipette tip (RPT, StarLab) and pipetted to mix. Samples were then incubated for 45 minutes at 22°C in the dark to complete reduction and alkylation. 4 µL of 1.25 M Azido-PEG3-Azide (diluted in DMSO) (Lumiprobe, #207) was added and incubated at 55°C for 20 minutes to quench TCEP. Samples were then approximately 100 µL in volume. 5 µL aliquots of each sample were taken and were diluted 5x in water and compared to a standard curve of BSA in 5x diluted lysis buffer using a microplate BCA assay (ThermoFisher, #23227). 25 µg of each sample was taken forward and the volumes of each lysate equalized using 50 mM TEAB, 5% (w/v) SDS.

For MS-AP experiments, eluates from affinity purification were brought to ∼5% (w/v) SDS and reduced and alkylated with 5 mM TCEP and 20 mM chloroacetamide for 45 minutes at 22°C in the dark before proceeding with digestion.

For the SILAC experiment, equal numbers of cells from each label were pooled prior to lysis; lysis and BCA assay were conducted as above. 500 µg of material was taken forward for digestion.

#### S-trap digestion

To each sample a 10% volume of 27.5% (v/v) phosphoric acid was added to acidify samples to ∼pH 2, completing denaturation. 6x volumes of Wash Buffer (100 mM HEPES, pH 7.55, 90% (v/v) methanol) was then added and the resulting solution was loaded onto a µS-trap (Protifi, #C02-Micro) using a positive pressure manifold (PPM) (Tecan M10), not more than 150 µL of sample at a time (∼80 PSI). In-house fabricated adaptors were used to permit the use of S-traps with the manifold. The S-traps were then washed three times with 150 µL Wash Buffer. To remove any remaining Wash Buffer, S-traps were centrifuged at 4000 × g for 2 minutes. To each S-trap, 30 µL Digestion Solution was added. This was 50 mM TEAB, pH 8.5, 1.25 µg Trypsin/lysC mix (Promega, V5072) for whole cell proteomics experiments and the amount of protease was reduced to 0.5 µg for MS-AP experiments. S-Traps were then loosely capped and placed in low adhesion 1.5 mL microfuge tubes in a ThermoMixer C (Eppendorf) with a heated lid and incubated at 37°C for 16 hours. Peptides were recovered by adding 25 µL Digestion Buffer to each trap and incubating at room temperature for 15 minutes before eluting by centrifugation at 400 × g. Traps were subsequently eluted with 40 µL 0.2% (v/v) formic acid (FA) and 25 µL 50% (v/v) Acetonitrile (ACN) in the same manner. Eluted samples were then dried in a vacuum centrifuge equipped with a cold trap.

For the SILAC experiment, 2x 250 µg aliquots of lysate were digested in parallel in S-Trap mini devices (Protifi, #C002-MINIX). Centrifugation was used in place of positive pressure to effect loading and washing of the traps, the wash volume was 350 µL. Digestion volume was 125 µL, elution volumes were 80 µL for each solvent and eluates were pooled from both digestions prior to drying under vacuum.

#### TMT labelling and clean-up

Samples were resuspended in 21 µL 100 mM TEAB pH 8.5. After allowing to come to room temperature, 0.5 µg TMTpro reagents (ThermoFisher, #A52045) were resuspended in 9 µL anhydrous ACN which was added to the respective samples and incubated at room temperature for 1 hour. A 3 µL aliquot of each sample was taken and pooled to check TMT labelling efficiency and equality of loading by LC-MS. Samples were stored at -80°C in the interim. After checking each sample was at least 96% TMT labelled, total reporter ion intensities were used to normalize the pooling of the remaining samples such that the final pool should be as close to a 1:1 ratio of total peptide content between samples as possible. This final pool was then dried in a vacuum centrifuge. The sample was acidified to a final 0.1% (v/v) Trifluoracetic Acid (TFA) (∼1000 µL volume) and FA was added until pH was <2. The sample was then cleaned up by solid phase extraction (SPE) using a 100 mg Strata C18-T SPE cartridge (Phenomenex, #8B-S004-EAK) and a PPM. The cartridge was wetted with 1 mL 100% (v/v) methanol followed by 1 mL ACN, equilibrated with 1 mL 0.1% TFA and the sample loaded slowly. The cartridge was washed twice with 1 mL 0.1% (v/v) TFA before being eluted with 250 µL 40% (v/v) ACN and 750 µL 80% (v/v) ACN. The samples were then dried under vacuum. TMT labels are shown below:

| TMT Label | ZFAND5/6 KO? | LPS (ng/ml) |
| --- | --- | --- |
| 126 | N | 0 |
| 127N | N | 0 |
| 127C | N | 0 |
| 129C | N | 100 |
| 130N | N | 100 |
| 130C | N | 100 |
| 131N | Y | 0 |
| 131C | Y | 0 |
| 132N | Y | 0 |
| 134N | Y | 100 |
| 134C | Y | 100 |
| 135 | Y | 100 |

SILAC samples were cleaned up in the same manner.

#### Basic pH reversed phase fractionation

Samples were resuspended in 40 µL 200 mM Ammonium formate (pH 10) and transferred to a glass HPLC vial. BpH-RP fractionation was conducted on an Ultimate 3000 UHPLC system (ThermoFisher) equipped with a 2.1 mm × 15 cm, 1.7µ Kinetex EVO column (Phenomenex, # 00F-4726-AN). Solvent A was 3% (v/v) ACN, Solvent B was 100% (v/v) ACN, and Solvent C was 200 mM ammonium formate (pH 10). Throughout the analysis Solvent C was kept at a constant 10%. The flow rate was 500 µL/min and UV was monitored at 280 nm. Samples were loaded in 90% Solvent A for 10 min before a gradient elution of 0–10% Solvent B over 10 min (curve 3), 10-34% Solvent B over 21 min (curve 5), 34-50% Solvent B over 5 mins (curve 5) followed by a 10-minute wash with 90% Solvent B. 15s (100 µL) fractions were collected throughout the run. Fractions containing peptide (as determined by A280) were recombined across the gradient to preserve orthogonality with on-line low pH RP separation. For example, fractions 1, 25, 49, 73, 97 are combined and dried in a vacuum centrifuge and stored at -20°C until LC-MS analysis. In this manner, 24 pooled fractions were generated.

For SILAC samples a slightly modified gradient of 0-32% B over 36 minutes was employed.

#### Mass spectrometry

Samples were analyzed on an Orbitrap Fusion instrument on-line with an Ultimate 3000 RSLC nano UHPLC system (ThermoFisher). Samples were resuspended in 10 µL 5% (v/v) DMSO/1% (v/v) TFA. 5 µL of each fraction was injected. Trapping solvent was 0.1% (v/v) TFA, Analytical solvent A was 0.1% (v/v) FA, Solvent B was ACN with 0.1% (v/v) FA. Samples were loaded onto a trapping column (300 µm x 5 mm PepMap cartridge trap (ThermoFisher, #174500)) at 10µL/minute for 5 minutes. Samples were then separated on a 75 cm x 75 µm i.d. 2 µm particle size PepMapNeo EasySpray C18 column (ThermoFisher, ES75750PN). The gradient was 3-10% Solvent B over 10 minutes, 10-35% Solvent B over 155 minutes, 35-45% Solvent B over 9 minutes followed by a wash at 95% Solvent B for 5 minutes and re-equilibration at 3% Solvent B. Eluted peptides were introduced by electrospray to the MS by applying 1.6 kV to the integrated needle. The MS was operated in SPS mode where MS1 Scans are obtained in the Orbitrap, CID-MS2 Scans are acquired in the Ion Trap and SPS-HCD-MS3 is acquired in the Orbitrap to resolve reporter ions.

For SILAC samples, the MS was operated in OT-IT mode with MS1 scans obtained in the Orbitrap (at 240’000 resolution) and MS2 scans obtained in the ion trap.

#### Data processing

Experiments were processed with Peaks, v12 (Bioinfor). Raw files were searched against the SwissProt Mouse Database with appended common contaminants, PSM FDR was controlled at 0.1% by decoy database search. For the ZFAND5/6 KO experiment, identified proteins and their abundances were output to .csv, imported to R and submitted to statistical analysis using LIMMA, a moderated t-test available through the Bioconductor package. LIMMA p-values were corrected for multiple hypothesis testing using the Benjamini-Hochberg method to generate an FDR (q-value) for each comparison. The MS-AP experiment was similarly processed but without statistical analysis. The mass spectrometry proteomics data have been deposited to the ProteomeXchange Consortium via the PRIDE partner repository with the dataset identifiers PXD071432 and 10.6019/PXD071432.

---

**Figure S1.**
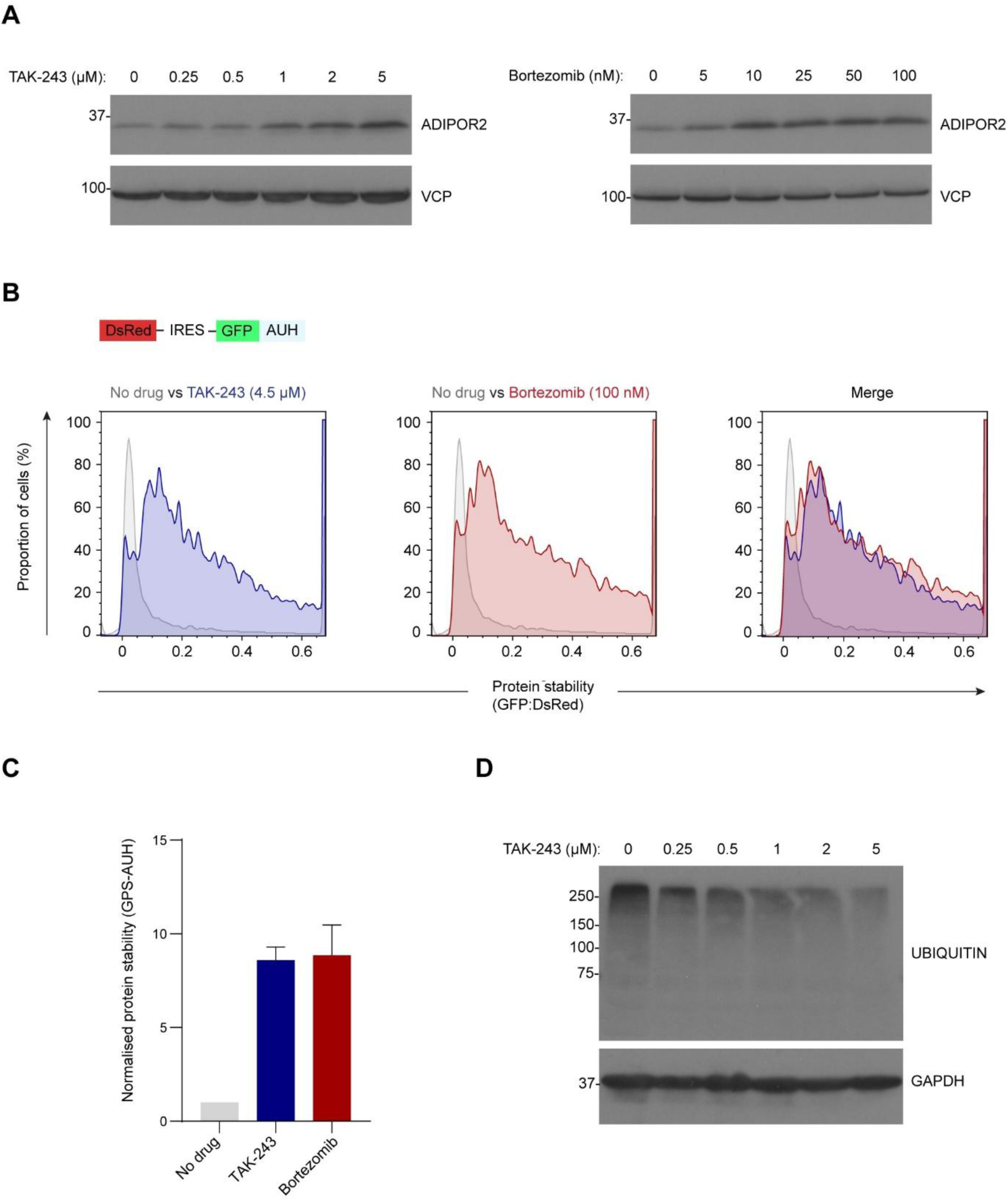
Optimising the TAK-243 and Bortezomib treatment conditions. **(A)** Initial optimisation of the TAK-243 and Bortezomib treatment conditions. The objective was to identify the concentration of each inhibitor required to maximally stabilise ADIPOR2, an unstable protein with a known E3 ubiquitin ligase in HeLa cells (Volkmar *et al*, 2022). TAK-243 inhibits global ubiquitination, while Bortezomib inhibits proteasome activity. The cells were treated with the two drugs for 17 hours. **(B)** Validation of the optimised drug treatment conditions in a flow cytometry-based protein stability assay. AUH is an unstable protein that is degraded in a conventional, ubiquitin-dependent fashion. The AUH coding sequence was cloned into a ‘GPS’ protein stability reporter plasmid (Yen *et al*., 2008). The plasmid encodes two fluorescent proteins (DsRed and GFP) upstream of its multiple cloning site. These two coding sequences are separated by an internal ribosome entry site (IRES) such that, upon translation, a single mRNA will encode two proteins: a DsRed protein, and the protein of interest fused to an N-terminal GFP tag. The DsRed protein is an internal control that reports on transcription and translation efficiency, and the GFP:DsRed ratio is a direct measure of protein stability. HeLa cells stably overexpressing the GPS-AUH plasmid were treated with TAK-243 or Bortezomib for 17 hours, and AUH stability was determined by flow cytometry. **(C)** Quantification of the assay shown in (B). The experiment was performed twice, and the mean protein stability values were normalised against the ‘no drug’ control. **(D)** Determining the effects of the TAK-243 on total cellular ubiquitination. Cells were treated with a TAK-243 titration for 17 hours and global ubiquitin conjugation was assayed via immunoblot.

**Figure S2.**
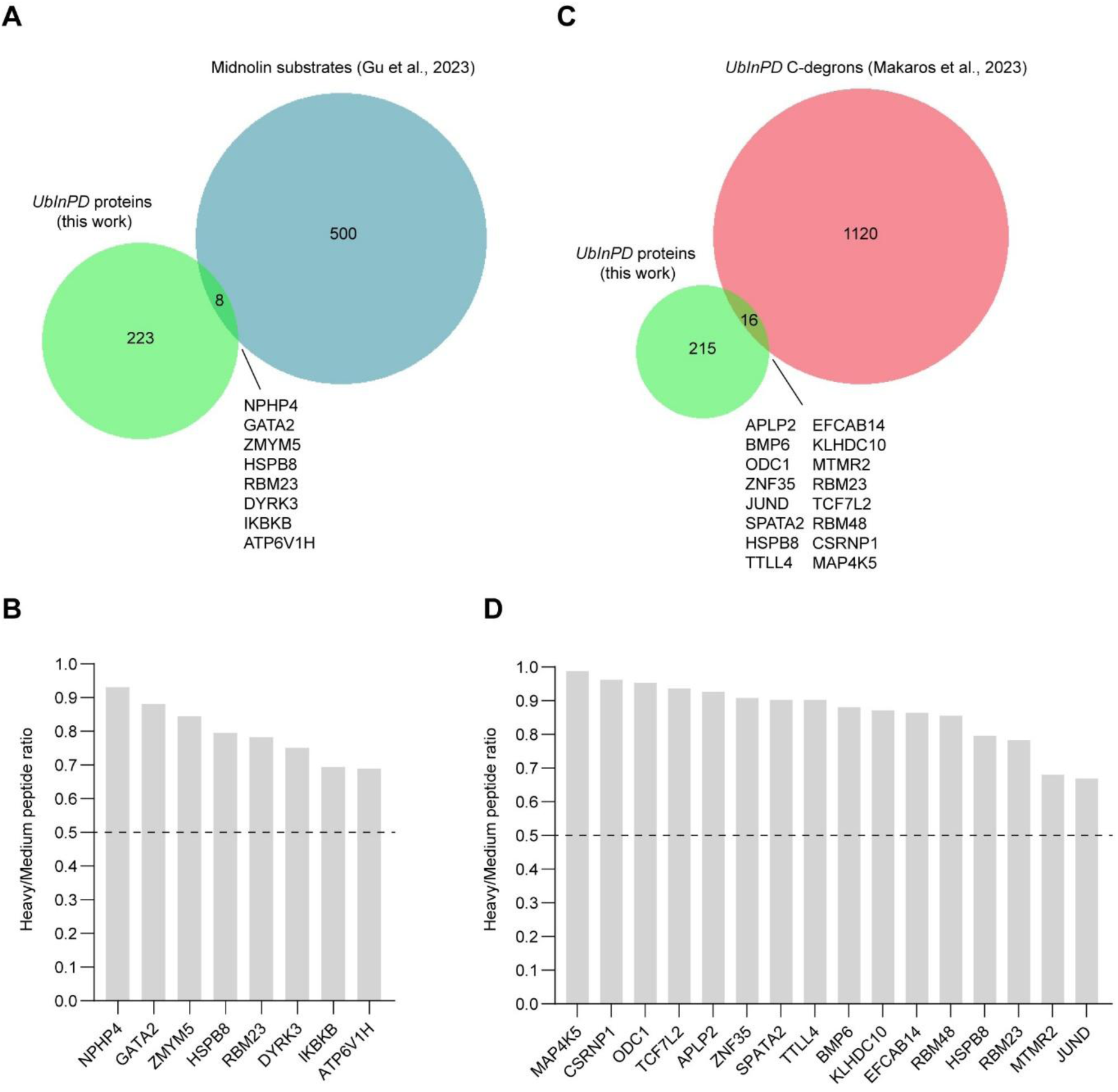
Identification of candidate Midnolin substrates and proteins bearing putative *UbInPD* C-degrons within the pulsed-SILAC screen. **(A)** Venn diagram showing the overlap between *UbInPD* proteins identified in this study and hits from a genetic screen for Midnolin substrates (Gu *et al*., 2023). A protein was considered a Midnolin substrate if its protein stability index (PSI) decreased by at least 50% when Midnolin was overexpressed. **(B)** Plot showing the H/M ratios of the 8 putative Midnolin substrates identified in this study. Any H/M ratio above the dashed line is positive; any ratio below it is negative. **(C)** Venn diagram showing the overlap between *UbInPD* proteins identified in this study and hits from a genetic screen for *UbInPD* C-degrons (Makaros *et al*., 2023). **(D)** Plot showing the H/M ratios of the 16 proteins that were hits in both this study and in the C-degron study.

**Figure S3.**
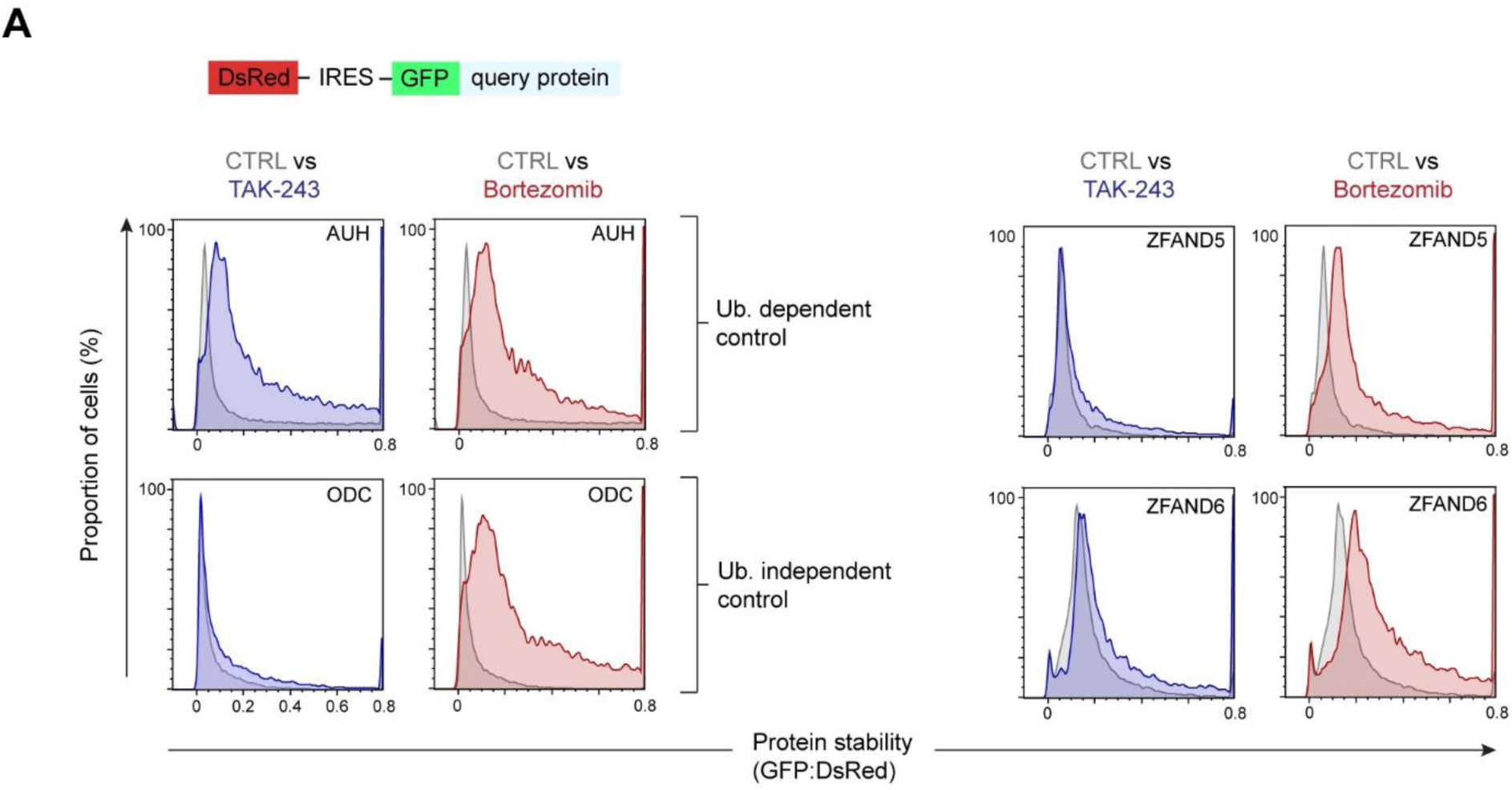
Validation of ZFAND5 and ZFAND6 by GPS assay. **(A)** ZFAND5, ZFAND6, AUH, and ODC were cloned into the GPS protein stability reporter plasmid. HeLa cells stably overexpressing each plasmid were treated with TAK-243 or Bortezomib for 17 hours and analysed via flow cytometry. The GFP/DsRed ratio is a direct measure of protein stability. AUH and ODC are ubiquitin-dependent and independent controls, respectively.

**Figure S4.**
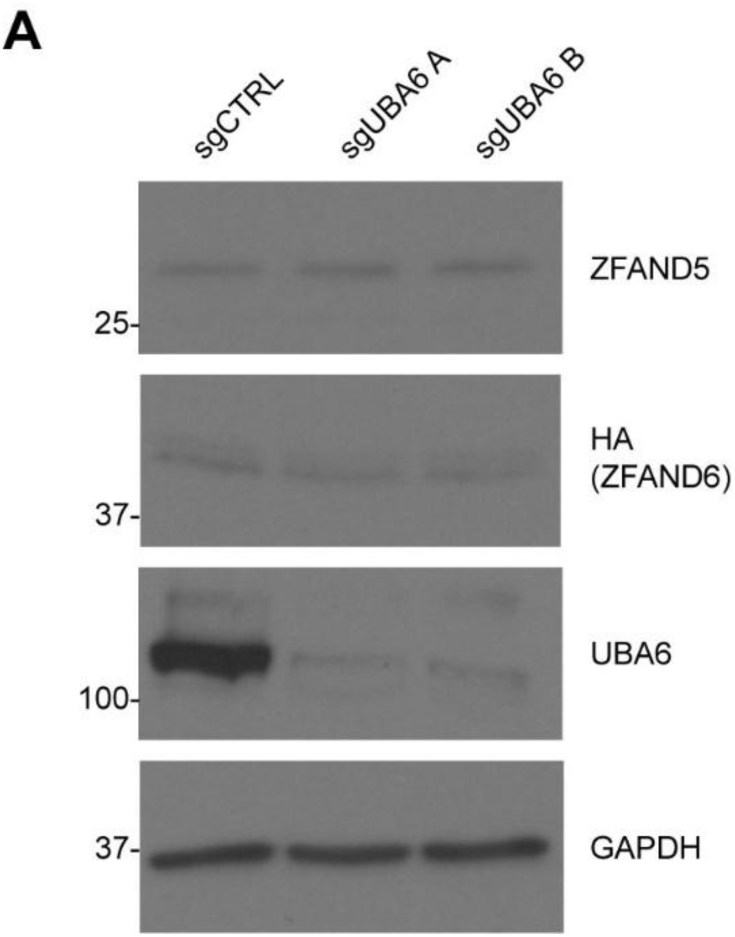
Analysis of ZFAND5 and ZFAND6 abundance in UBA6-deficient cells. **(A)** The minor E1 ubiquitin activating enzyme UBA6 was deleted via CRISPR-KO in cells stably overexpressing HA-ZFAND6. Two independent sgRNAs were used.

**Figure S5.**
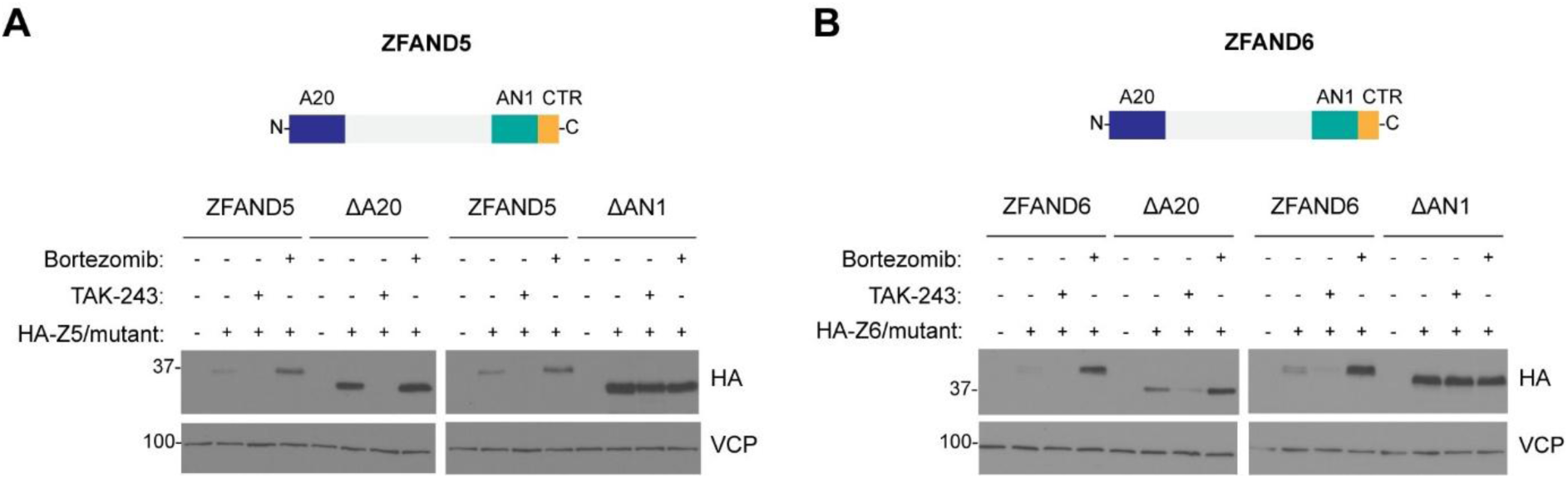
Characterisation of the domains required for ZFAND5/6’s ubiquitin-independent proteasomal degradation. **(A)** Cells stably overexpressing 3xHA ZFAND5 or indicated truncation mutants (denoted by the Δ symbol) were treated with TAK-243 or Bortezomib for 17 hours. **(B)** As in (A) but with 3xHA ZFAND6.

**Figure S6.**
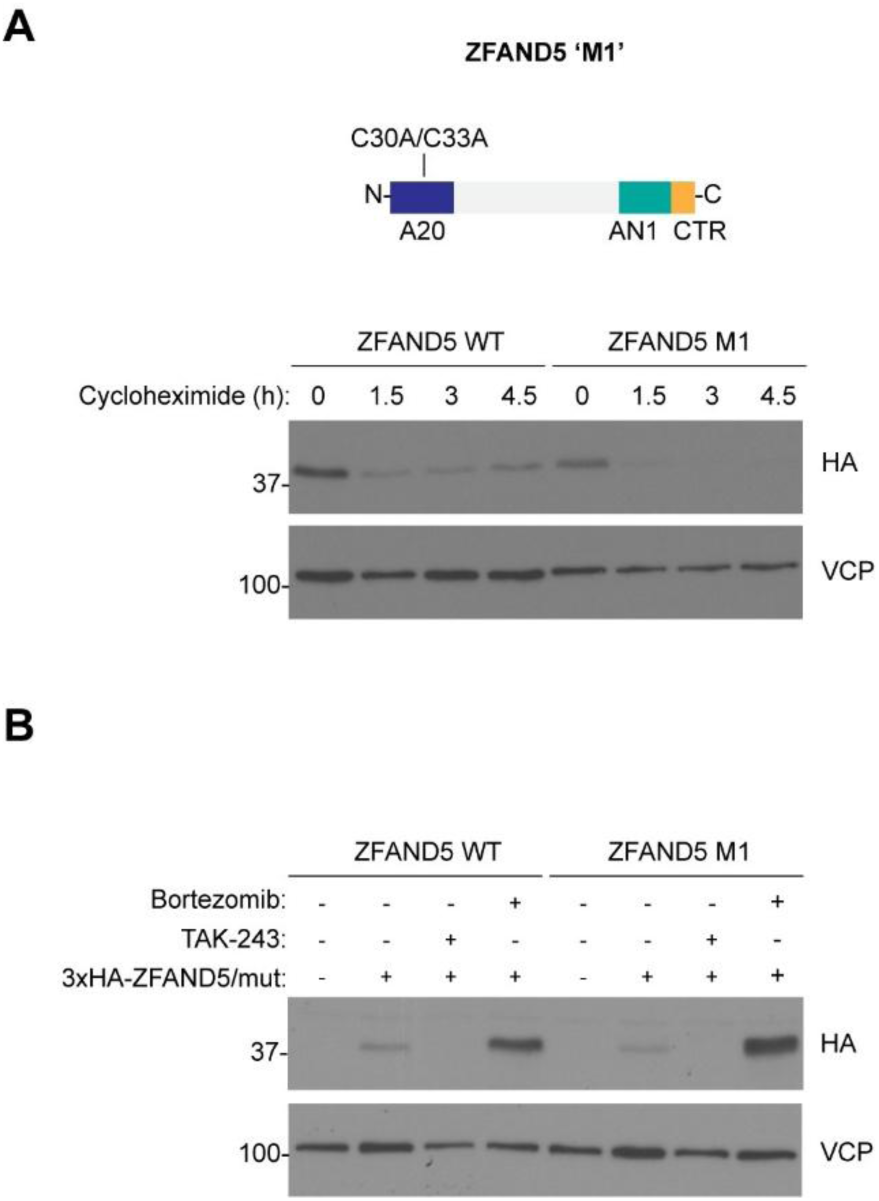
Point mutations in the ZFAND5^A20^ domain do not change ZFAND5’s unusual mode of degradation. **(A)** Cycloheximide chase assay profiling the stability of ZFAND5 or the M1 mutant. ZFAND5 M1 has two point mutations (C30A/C33A) in its A20 domain which completely abolish its ability to bind to ubiquitinated proteins (Lee *et al*., 2018). Cells stably overexpressing the two proteins were treated with 10 µg/ml cycloheximide as indicated. The two proteins carried 3xHA tags at their N-termini. **(B**) The cells from (A) were treated with TAK-243 or Bortezomib for 17 hours. Importantly, the ZFAND5 WT panel (lanes 2-4) is also shown in (Fig. 2B). Mut: Mutant (M1).

**Figure S7.**
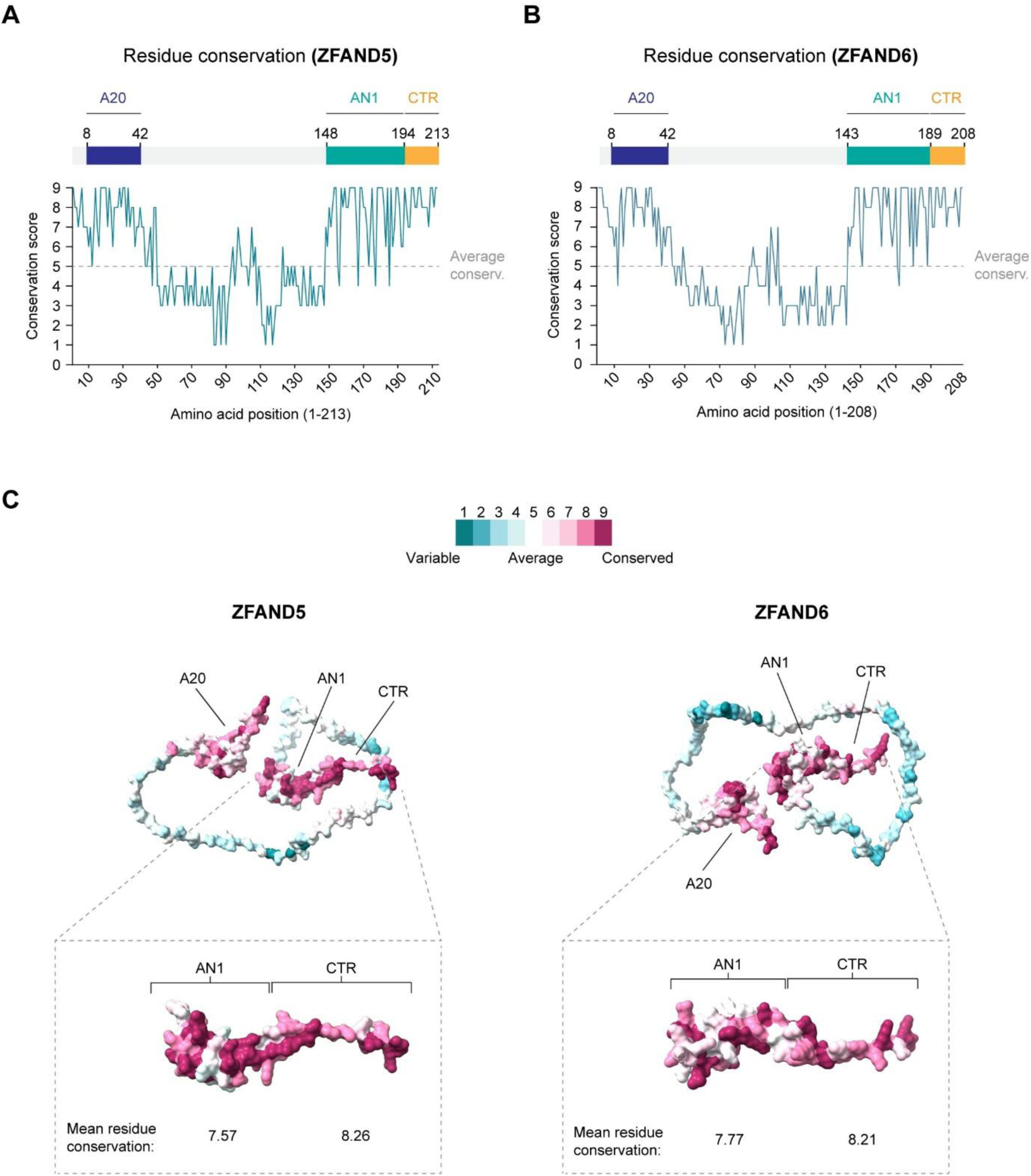
The ZFAND5/6 degrons are deeply conserved. **(A)** Plot showing the conservation of residues across ZFAND5 (according to the CONSURF algorithm, which measures residue conservation across different organisms). Scores range from 0 (highly variable) to 9 (highly conserved). **(B)** As in (A) but for ZFAND6. **(C)** Alphafold structures of ZFAND5/6 coloured by residue conservation. The AN1/CTR degrons are magnified and shown in panels beneath the main structures. The numbers beneath the domains indicate the mean conservation of residues within the AN1 and CTR domains.

**Figure S8.**
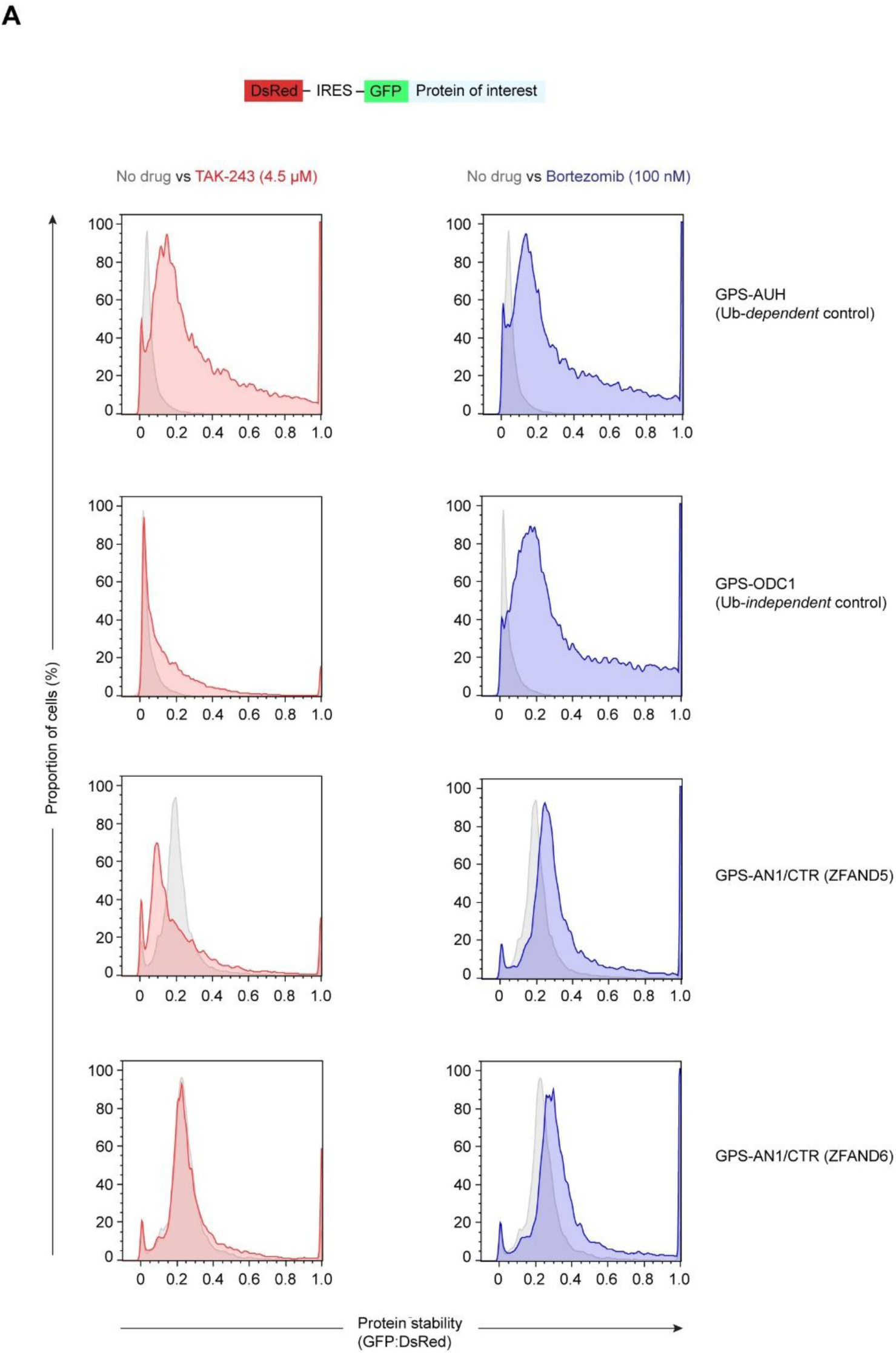
The ZFAND5/6 degrons are portable. **(A)** The AN1/CTR degrons were cloned into the GPS reporter plasmid. Cells stably expressing the two reporters, or the indicated controls, were treated with TAK-243 or Bortezomib for 17 hours.

**Figure S9.**
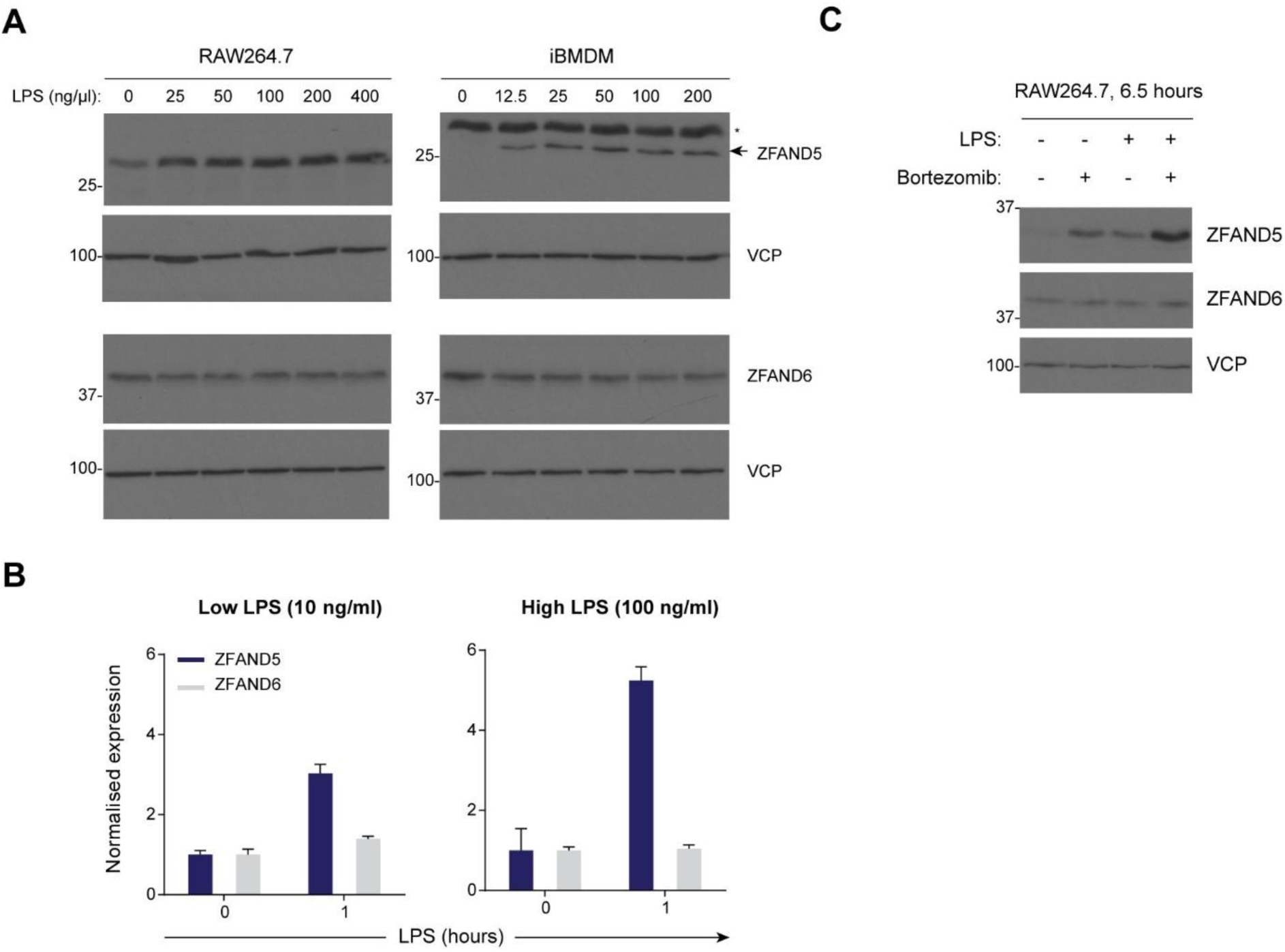
ZFAND5 is upregulated in LPS-treated mouse macrophages. **(A)** ZFAND5 is upregulated at the protein level following LPS stimulation. RAW264.7 cells (left) or iBMDMs (right) were treated with the indicated LPS titrations for 17 hours and the samples were resolved via immunoblotting. The asterisk (*) indicates a non-specific band. **(B)** Rapid transcriptional upregulation of ZFAND5 following LPS stimulation. RAW264.7 cells were treated with LPS (10 ng/ml or 100 ng/ml, as indicated) for 1 hour and ZFAND5/6 expression was determined by RT-qPCR. Expression was normalised to GAPDH and presented as fold change relative to the untreated control. Error bars: mean ± SD from three technical replicates. **(C)** ZFAND5 is upregulated following LPS stimulation and is rapidly degraded by the proteasome. RAW264.7 cells were treated with LPS (10 ng/ml) or Bortezomib (100 nM) singly and in combination for 6.5 hours, and ZFAND5 protein levels were determined by immunoblotting.

**Figure S10.**
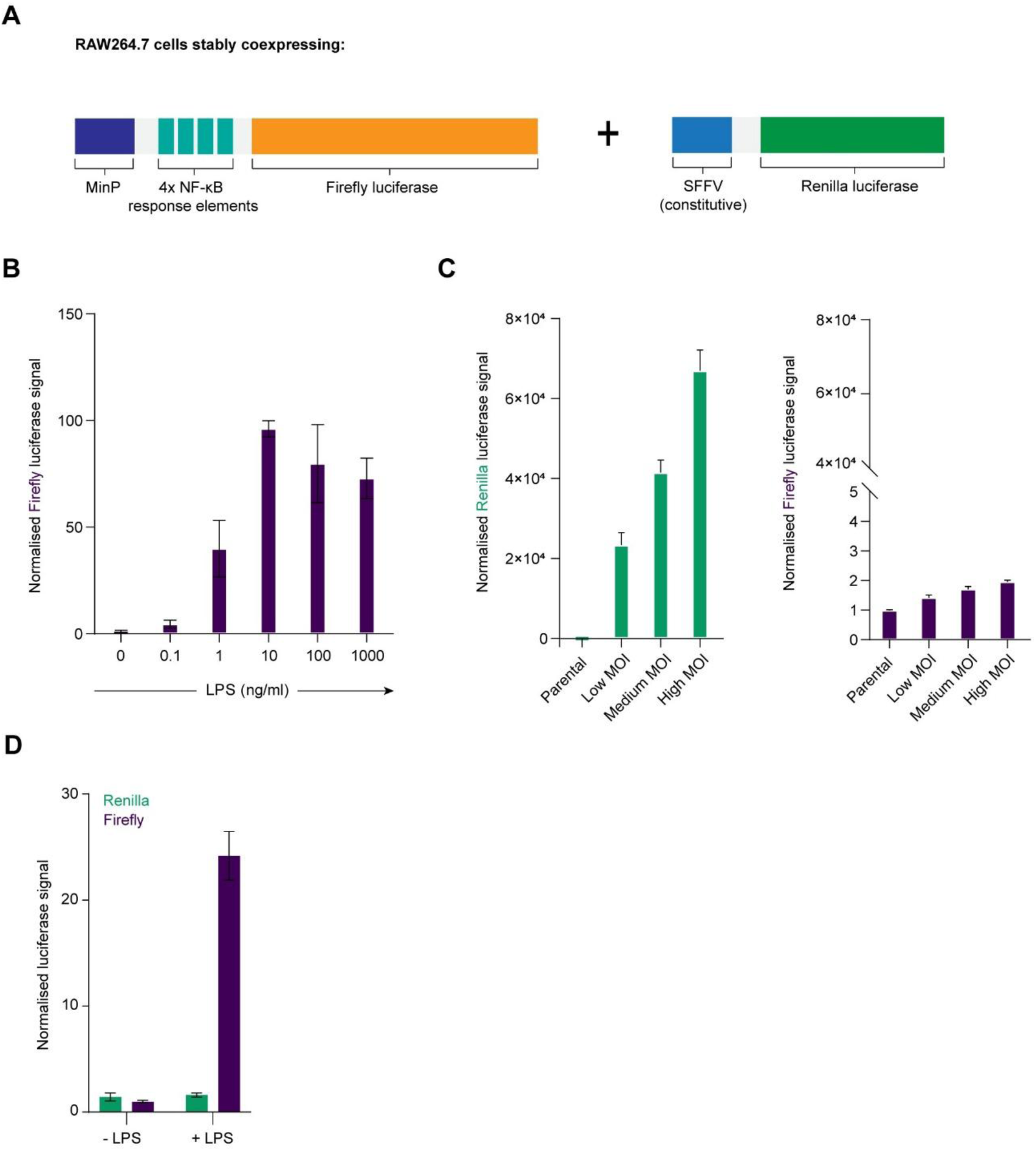
Engineering a cell line that reports on NF-κB activation in the TLR4 pathway. **(A)** Schematic representation of the reporter cell line. MinP: Minimal promoter. SFFV: Spleen focus-forming virus, a strong constitutive promoter. Cells expressing just the Firefly luciferase cassette (the ‘parental cell line’) were obtained and analysed in (B) and transduced with the Renilla cassette as will be described. **(B)** Rapid induction of Firefly luciferase signal in the parental cell line. Cells were treated with a 10-fold serial dilution of LPS for 5 hours. Firefly luciferase signal was expressed as fold change relative to the untreated control. Error bars: mean ± SD from three technical replicates. **(C-D)** Expression of Renilla luciferase in the engineered cell line. Cells were engineered to express constitutive Renilla to serve as a control for subtle differences in cell seeding density and cell lysis efficiency. The parental cell line was transduced with a titration of virus carrying the Renilla cassette, and Firefly (right) and Renilla (left) signal was assayed sequentially in untreated cells (C), or in cells treated with LPS (10 ng/ml) for 5 hours (D). MOI: Multiplicity of infection. Renilla expression was robust and showed minimal cross-detection in the Firefly channel. Cells transduced at the ‘medium’ MOI were assayed in (D) and used in all subsequent experiments. Data were analysed as in (B).

**Figure S11.**
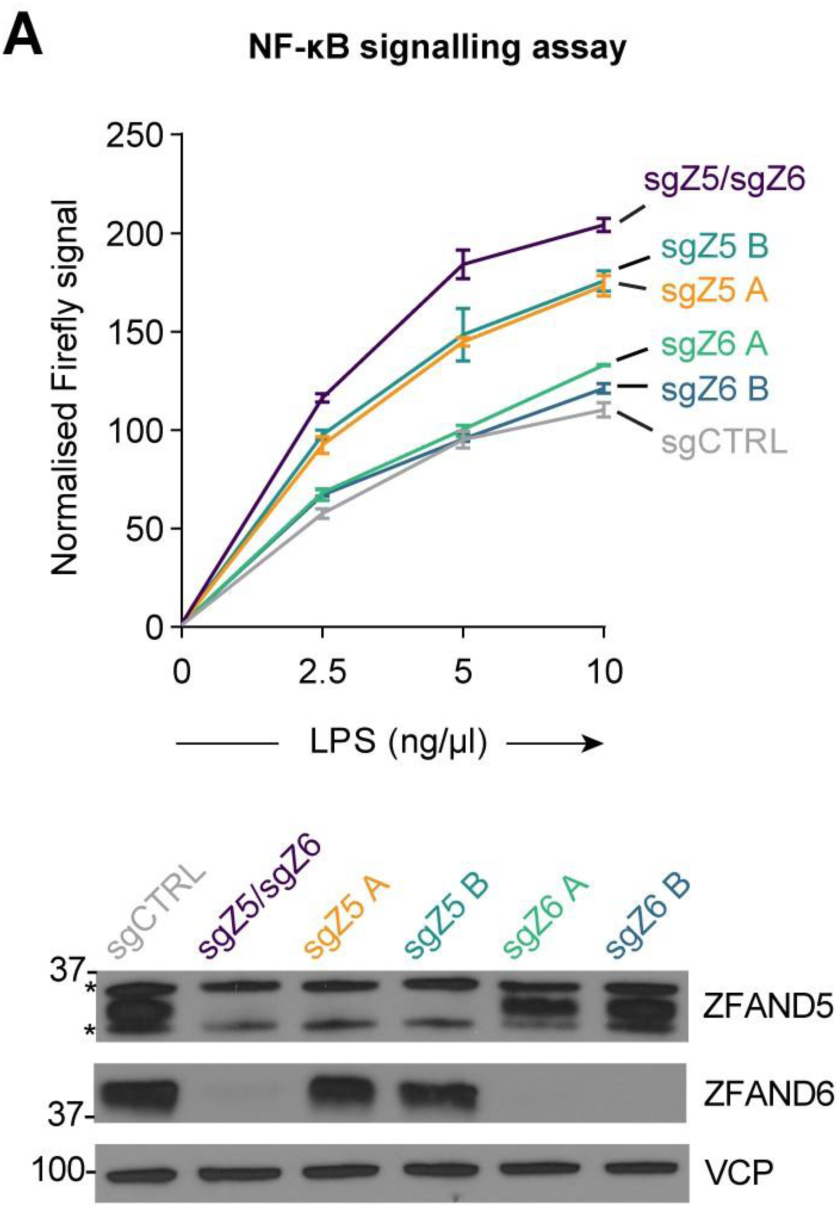
ZFAND5/6 restrain NF-κB activation in the TLR4 pathway. **(A)** ZFAND5 and ZFAND6 restrain NF-κB activation. Single ZFAND5 and ZFAND6 knockout cells were compared with double ZFAND5/6 knockout cells. Knockout efficiency was assayed via immunoblot. Asterisks (*) indicate non-specific bands. Cells were treated with LPS as indicated for 6 hours. Firefly signal was normalised against Renilla signal and presented as fold change relative to the untreated, non-targeting sgRNA control (sgCTRL). Error bars: mean ± SD from three technical replicates.

**Figure S12.**
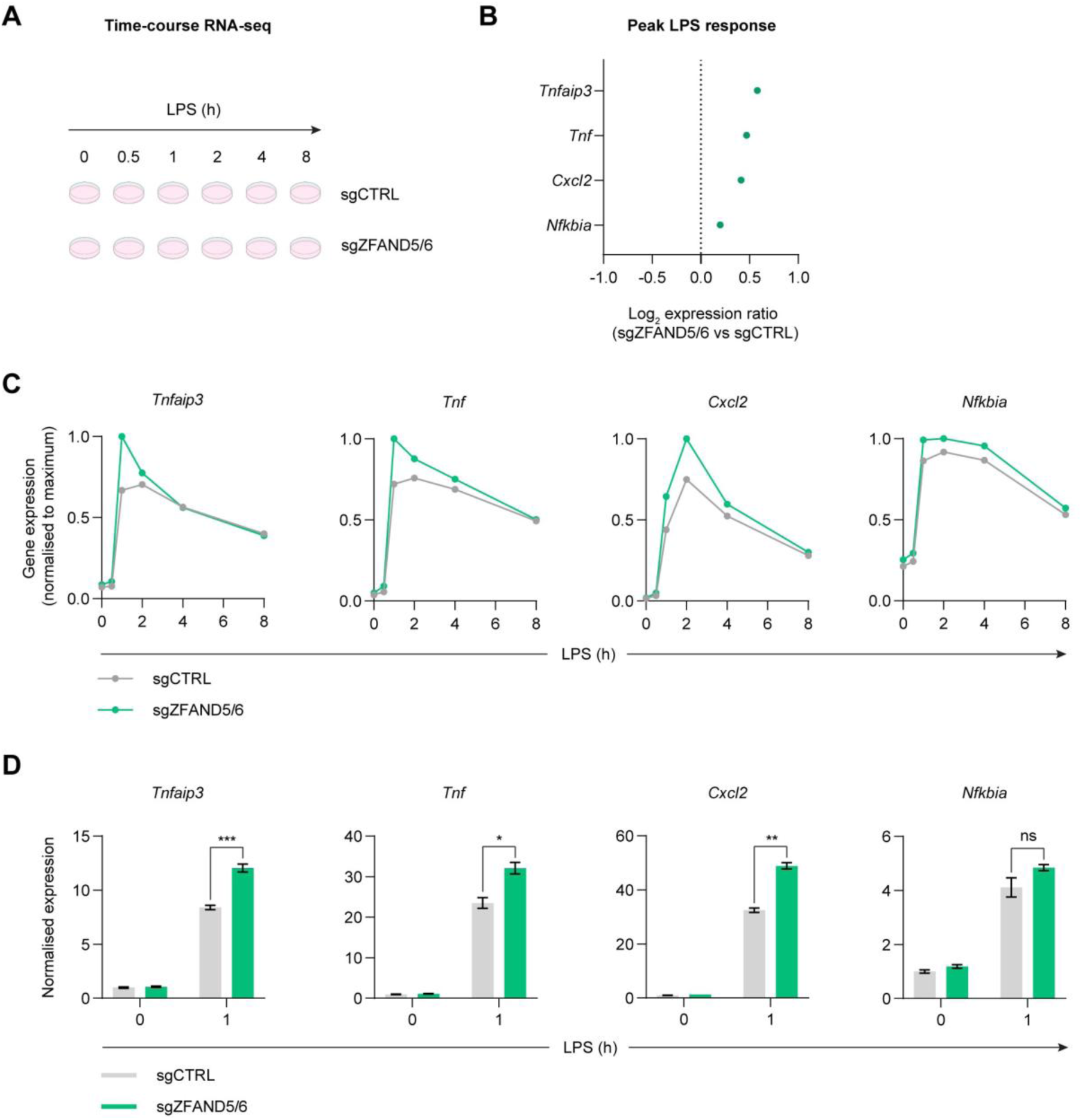
RNA-seq analysis reveals enhanced expression of early NF-κB response genes in LPS-treated ZFAND5/6 knockout cells. **(A)** Outline of the RNA-seq experiment. Control (sgCTRL) or ZFAND5/6-deficient (sgZFAND5/6) cell populations were treated with LPS (1 ng/ml) across an 8-hour time-course. **(B)** Comparative expression of four canonical NF-κB response genes between sgCTRL and sgZFAND5/6 cells. Because TLR4 signals via multiple transcription factors that exhibit significant crosstalk (NF-κB, AP1, CREB, and IRF3) (Duan *et al*, 2022), it was necessary to identify genes that are specifically or primarily regulated by NF-κB. Candidate genes were selected from a published list of LPS-responsive genes that are directly regulated by NF-κB (Sharif *et al*, 2007), filtered for expression in RAW264.7 cells, and restricted to genes exhibiting maximal induction within 2 h of LPS stimulation to minimise crosstalk effects. This yielded a panel of four early NF-κB response genes (*Tnf*, *Nfkbia*, *Cxcl2*, and *Tnfaip3*). For each gene, expression was compared at the time of maximal induction following LPS stimulation. Values are shown as log₂(sgZFAND5/6/sgCTRL). **(C)** Temporal expression profiles of the early NF-κB response genes following LPS stimulation. Expression values were normalised to the maximum value observed for each gene to permit comparison on a common scale. **(D)** Validation of the RNA-seq results by RT-qPCR. Expression of *Tnfaip3*, *Tnf*, *Cxcl2* and *Nfkbia* was quantified following LPS stimulation (1 ng/ml; 0 or 1 hour). Transcript abundance was normalised to GAPDH and expressed relative to the untreated, sgCTRL samples. Data are shown as mean ± SD. Statistical significance was determined using a two-tailed paired Student’s t-test. Asterisks indicate significance: *P < 0.05, **P < 0.01, ***P < 0.001. Ns: not significant.

**Figure S13.**
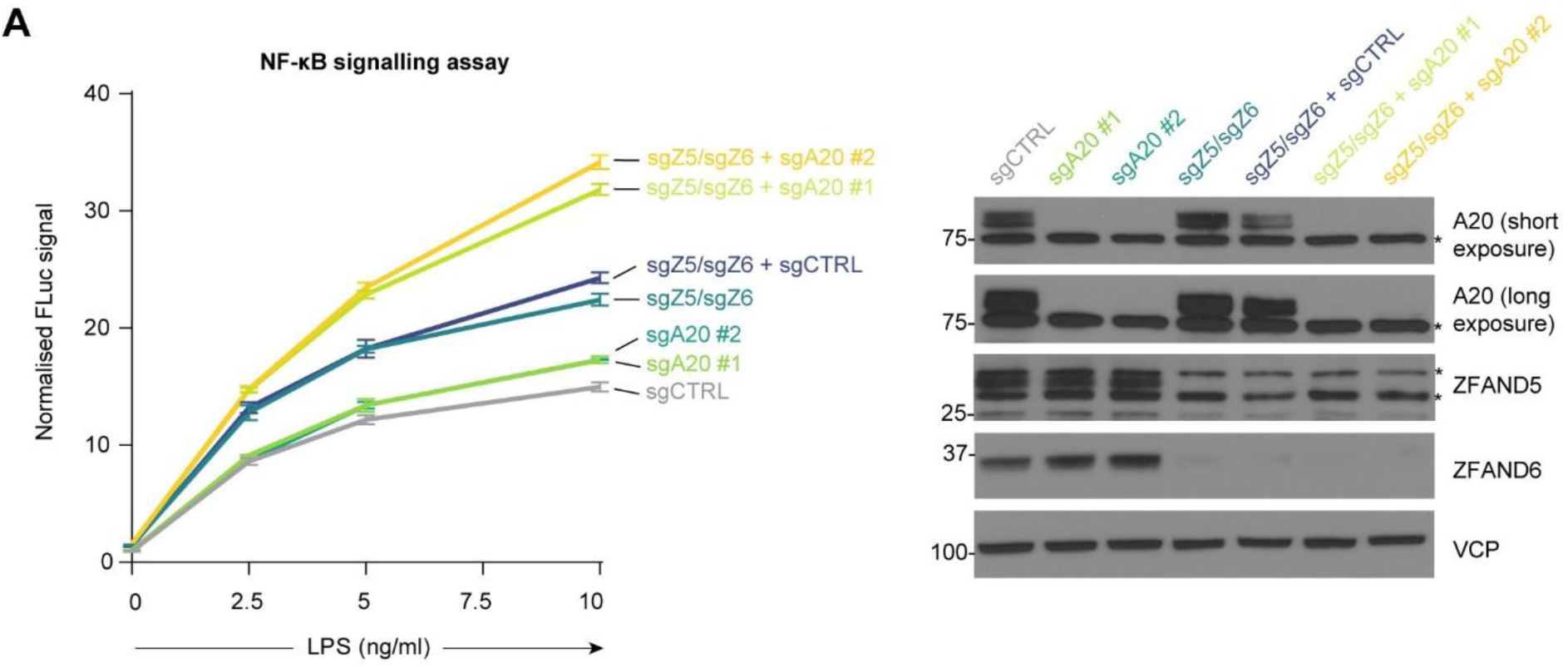
Genetic analysis of ZFAND5/6 and A20. **(A)** There is a strong genetic interaction between ZFAND5/6 and A20. A20 was deleted in the reporter cell line via two independent sgRNAs, either singly or in the ZFAND5/6 double knockout background. Knockout efficiency was assessed by immunoblotting. Asterisks (*) indicate non-specific bands. Cells were treated with LPS as indicated for 6 hours. Firefly signal was normalised against Renilla signal and presented as fold change relative to the untreated, non-targeting sgRNA control (sgCTRL). Error bars: mean ± SD from three technical replicates.

**Figure S14.**
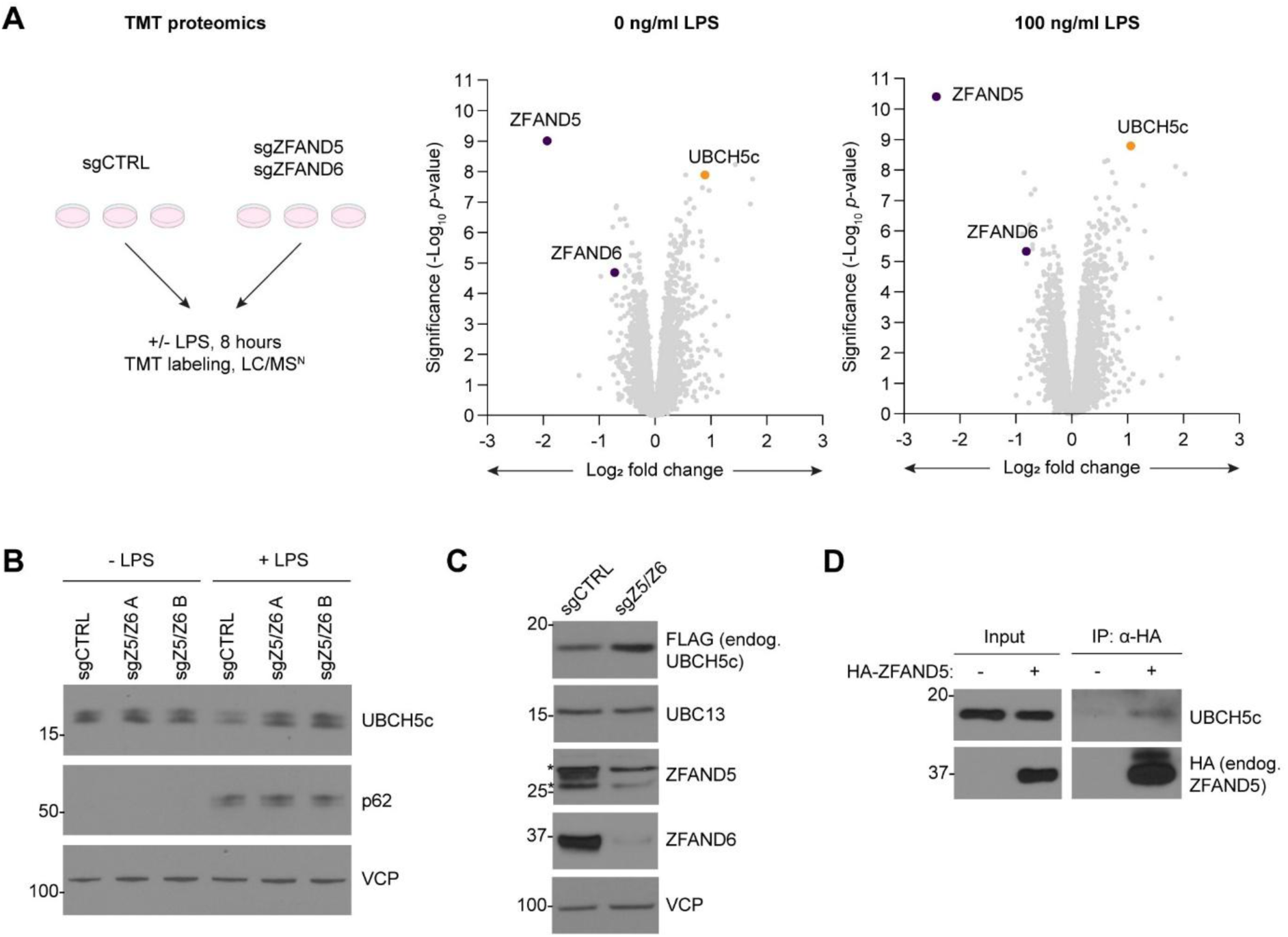
ZFAND5/6 regulate UBCH5c abundance. **(A)** Identification of candidate ZFAND5/6 substrates via quantitative proteomics. ZFAND5/6 knockout or control cells (populations, not clones) were treated ± LPS (100 ng/ml) for 8 hours. UBCH5c was significantly upregulated in both the untreated and treated ZFAND5/6 knockout cells (left and right panels, respectively). **(B)** UBCH5c degradation is blocked in ZFAND5/6 knockout cells. Cells were treated with LPS (10 ng/ml) overnight. The UBCH5c antibody may cross-react with the UBCH5c paralogues, UBCH5a and UBCH5b. **(C)** UBCH5c is upregulated in ZFAND5/6 knockout cells at steady state. ZFAND5/6 were knocked out in cells expressing UBCH5c-FLAG (endogenously tagged). UBC13 was included as a control E2. Asterisks (*) indicate non-specific bands. **(D)** ZFAND5 and UBCH5c physically associate. HA-ZFAND5 (endogenously tagged) was immunoprecipitated under non-denaturing conditions. The cells were pretreated with LPS (10 ng/ml) and Bortezomib (100 nM) for 6 hours before the IP. (-) Wild-type cells; (+) cells expressing endogenously tagged HA-ZFAND5.

**Figure S15.**
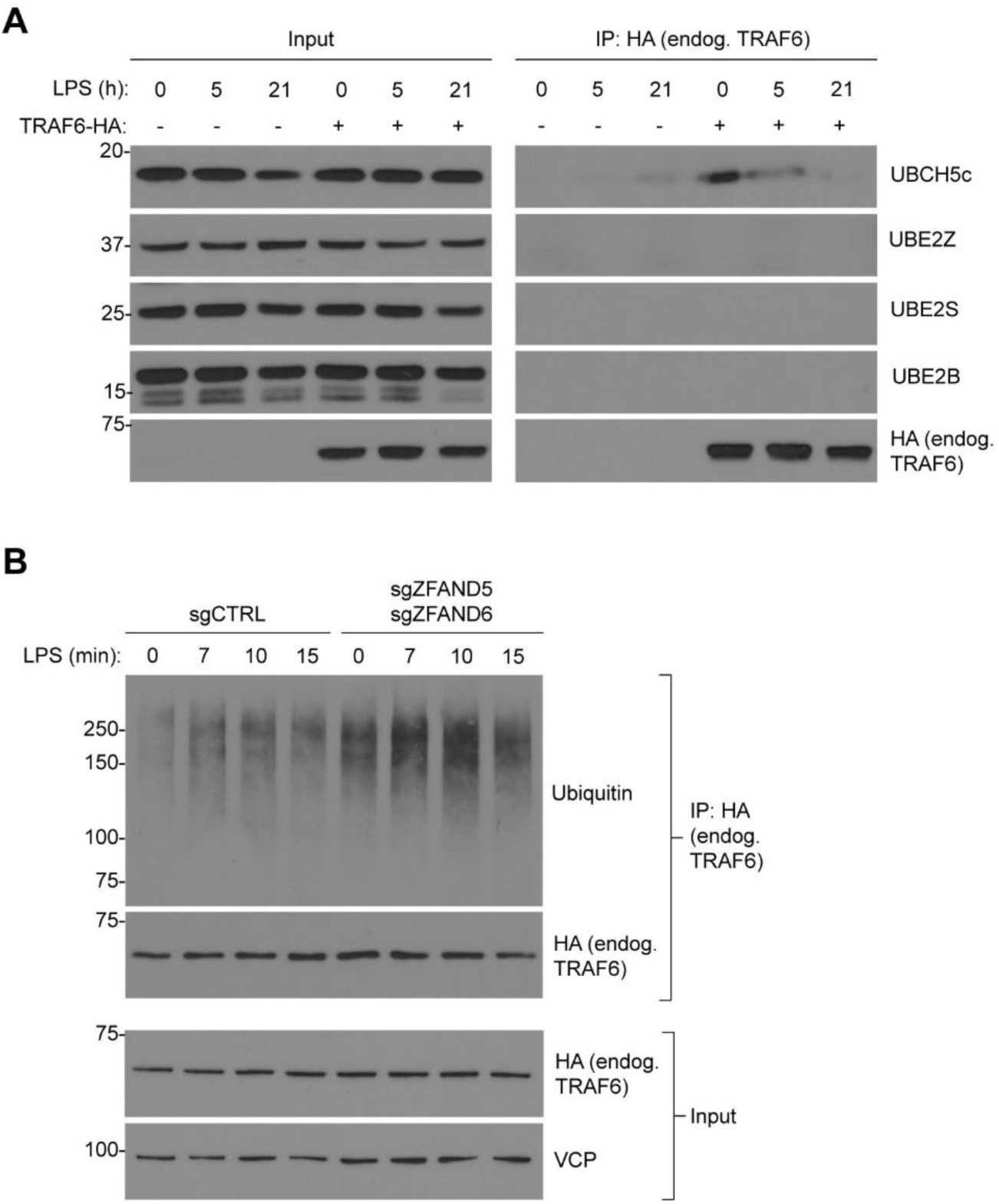
ZFAND5/6 and p62 suppress TRAF6 activity. **(A)** TRAF6 associates with UBCH5c, and this interaction is lost with sustained LPS stimulation. Wild-type cells, or cells expressing TRAF6-HA (endogenously tagged) were treated with LPS (10 ng/ml) across a 0-, 5-, and 21-hour LPS treatment time-course. UBE2Z, UBE2S, and UBE2B were included as control E2s. (-) Wild-type cells; (+) TRAF6-HA cells. **(B)** TRAF6 activity is heightened in ZFAND5/6 knockout cells. ZFAND5/6 were knocked out in cells expressing TRAF6-HA (endogenously tagged) and TRAF6 was immunoprecipitated under denaturing conditions across a short time-course.

